# Beyond Recombination: Exploring the Impact of Meiotic Frequency on Genome-wide Genetic Diversity

**DOI:** 10.1101/2025.03.11.642586

**Authors:** Louis Ollivier, Brian Charlesworth, Fanny Pouyet

## Abstract

An important aim of population genetics is to elucidate the processes affecting genetic diversity across regions of the genome and across species. Canonical population genetic models of sexually reproducing species define the rate of meiotic recombination in terms of the frequency of recombination events per site per sexual generation. This paper studies the interplay of several factors with respect to their effects on neutral genetic diversity in a facultatively sexual, diploid, unicellular species such as yeast. The relevant factors are the prevalence of meiosis versus mitosis, the recombination rate, and the selection and dominance coefficients at loci under positive selection. We assume that many generations of mitotic cell divisions are interspersed with episodes of sexual reproduction, in which all individuals in the population undergo meiosis, followed by random matings among the resulting gametes. Our findings reveal that a single hard selective sweep can reduce neutral nucleotide site diversity across the entire genome, provided that the frequency of meiotic events is sufficiently low, and that the effects of a selective sweep on levels of neutral diversity at sites linked to the target of selection can be markedly different from those predicted by standard models of sweeps based on obligate sexuality. Species that reproduce by facultative sex are thus likely to exhibit unusual patterns of genetic diversity.

**Author summary:** In this study, we explored how different sexual strategies influence genetic diversity. Specifically, we looked at how the balance between sexual reproduction (which involves meiosis) and asexual reproduction (which involves mitosis) affect genetic variation in a species like yeast. Our research focused on the effects of recombination rates and the role of selective sweeps — when a beneficial genetic variant spreads rapidly through a population — on genetic diversity. We found that, in species with facultative sex, a selective sweep can dramatically reduce genetic diversity across the genome, but only if sexual reproduction occurs infrequently. We compared our findings to existing models and also developed a new mathematical framework for understanding the effects of sweeps when sexual reproduction is rare. The fact that facultative sex can reduce genetic diversity over the entire genome is likely to complicate inferences about population size and evolutionary dynamics in species with mixed reproductive strategies.

## Introduction

A major goal of population genetics is to identify and quantify the mechanisms that shape genetic variation, by comparing data on DNA sequence variation in natural populations with theoretical predictions, including computer simulations [1]. In addition to models based on the coalescent process [2,3], researchers also use forward-in-time simulations such as SLiM [4] to study complex evolutionary scenarios, including polygenic selection (see, for example, [5,6]) and intricate demographies [7]. All such population genetics models incorporate essential parameters such as mutation rates, population sizes, selection coefficients, and recombination rates [8,9].

In standard population genetics models, the rate of recombination is expressed in units of crossovers and gene conversion events per site per sexual generation. But the passage from one generation to the next is not achieved through meiosis alone, especially in facultatively sexual organisms such as unicellular eukaryotes or simple multicellular species, where mitotic divisions may also contribute to reproduction. In rotifers, for example, some clones have permanently lost the ability to reproduce sexually due to variants like the *op* allele, while others remain as cyclical parthenogens that switch to sex under specific cues such as high population density [10]. Sexual episodes often occur near population peaks, supporting the idea that sex is an anticipatory strategy rather than a response to stress [11]. Similar reproductive strategies are observed in aphids, green algae, fungal pathogens, and yeasts, where long clonal phases alternate with rare meiotic events. Asexual modes of reproduction with modified forms of meiosis such as automixis, may retain aspects of recombination or segregation, but typically do not involve genetic exchange between individuals. Consequently, the effective recombination rate, a factor modulating the effects of selection at linked sites, can vary dramatically across taxa, depending on both the recombination rate per meiosis and the frequency of sexual reproduction. For instance, in the budding yeast *Saccharomyces cerevisiae*, cells can regulate their division strategy, and a sexual generation with meiotic recombination occurs only once every 500 to 10,000 generations in nature [12,13]. Studies on natural populations of its relative *S. paradoxus* suggest that the frequency of sexual reproduction in that species ranges between 1/10,000 and 1/1,000 per cell division [14,15].

Research on the population genetics of facultative sex remains limited compared to models of obligate sexual or asexual reproduction. For instance, neutral coalescent models of the effects of facultative sex have primarily focused on gene conversion between homologous chromosomes during the asexual phase as a key form of recombination affecting neutral genetic diversity. These models show that such gene conversion events can substantially reduce neutral genetic diversity in facultatively sexual species [16]. This counteracts the effect of long-term asexuality, which causes long coalescence times for a pair of alleles within the same individual, thereby increasing the net level of sequence divergence among pairs of alleles [17]. Despite these advances, current models of facultative sex [5,16,18] have largely overlooked the role of natural selection in shaping genetic diversity in this context, which is the focus of our paper.

To address this gap, we explore the interplay between the frequency of sex and other evolutionary forces within the framework of genetic hitchhiking, a term introduced by Maynard Smith and Haigh in 1974 in the context of the spread of a beneficial mutation [19]. More generally, hitchhiking refers to the reduction of genetic variation at neutral sites that occurs in the vicinity of a variant under selection. It can either be caused by a beneficial mutation that increases in frequency in a population (a selective sweep) or by linked deleterious mutations that decrease in frequency (background selection) [20]. Here we specifically focus on selective sweeps. The recombination rate is a core parameter affecting the strength of hitchhiking, due to the effect of recombination in breaking down associations between neutral and selected loci [9,19].

It is well established that recombination rates vary across the genome of a single individual, between individuals of the same species, and between related species – for a review, see [21]. Interestingly, it is still unclear why a limited range of genetic diversity is observed between species, even among those with different census population sizes (Lewontin’s paradox) [22–24]. In facultatively sexual species like yeast, the frequency of meiotic generations (where recombination events are frequent) versus mitotic generations (where they are rare or absent) must potentially reduce their levels of genetic diversity compared to obligately sexual species with the same per generation mutation rate and population size, due to the stronger genome-wide effects of hitchhiking.

To start to address this question, we have developed a simulation-based model of selective sweeps, which includes both the recombination rate per site per meiosis and the frequency of meioses relative to mitosis. We assume that a population of diploid cells reproduces mitotically, with all members of the population occasionally undergoing meiosis followed by random mating of the haploid gametes, *i.e* outcrossing; this event is followed by another cycle of mitotic cell divisions. This is intended to capture the properties of the budding yeast *Saccharomyces cerevisiae*, a well-characterized facultatively sexual species that undergoes sexual reproduction in stressful environments, such as under nutrient deprivation [25]. While other reproductive strategies, such as haploselfing, occur in *S. cerevisiae* [13], for simplicity this study concentrates on outcrossing and the clonal reproduction of diploid individuals, in order to provide a proof of concept. We use a simplified dichotomy where asexual reproduction is equated with mitotic divisions and sexual reproduction with meiosis followed by random mating. While this framework is appropriate for organisms such as *Saccharomyces cerevisiae*, which alternates between clonal (mitotic cell divisions) and sexual reproduction (meioses), we acknowledge that many forms of asexual reproduction in other taxa involve modified meioses, sometimes including recombination and segregation. The defining feature we focus on here is the absence of genetic exchange between individuals during asexual reproduction, rather than the precise cellular mechanism involved.

We chose *S. cerevisiae* because its evolutionary parameters, including mutation rates, have been estimated from laboratory experiments [26], and its high diversity in wild populations [27,28] suggests a large effective population size, which typically supports the efficiency of natural selection unless hitchhiking is pervasive. Interestingly, despite its large census and effective population size, natural *S. cerevisiae* isolates are often highly homozygous diploids (e.g., [12,28]), suggesting that there is a high level of inbreeding, which we show greatly enhances the effects described here (see section 4 of the S1 Appendix). Here, we test whether rare sex may also play a role in shaping diversity patterns, possibly through the widespread effects of hitchhiking. Indeed, we show that a low frequency of meioses can create a footprint of natural selection at a genome-wide level, and that its effects on neutral diversity are not necessarily accurately predicted by the standard models of selective sweeps. Our results suggest that including the frequency of meioses is essential for correctly modeling the genome-wide impact of selection in facultatively sexual organisms.

## Results

### A low meiotic frequency can cause a sweep to affect the entire genome

To explore how the frequency of sex and recombination influences genetic diversity after the completion of a selective sweep, we first investigated the interaction between selection and the frequency of sexual reproduction (*i.e*, meiosis), denoted by *α.* We compared a case with obligatory sex (*α* = 1) to one with facultative sex (α = 0.01, where the interval between two sexual generations was fixed at 1/*α* = 100 generations). We tested whether the meiotic frequency affects neutral genetic diversity (*π*) by first comparing a control case (the absence of selection) to a test case with a positively selected site in the middle of the simulated chromosome. The product of the frequency of recombination per basepair per meiosis (*ρ_α_*) and *α* is denoted by the parameter *ρ*. For purposes of comparisons among different values of *α*, *ρ* is kept constant here, this means that when α is small, *ρ_α_* is increased such that *ρ_α_* × α remains constant. It can be regarded as measuring the net rate of recombination per basepair over all generations, mitotic and meiotic. A single beneficial mutation with selection coefficient *s* and dominance coefficient *h* is introduced into a population of *N* diploid individuals and allowed to proceed to fixation, such that all individuals in the population are homozygous for the mutation (see Simulation Methods in the Material and Methods for full details).

In the absence of selection, the nucleotide site diversity across the genome was similar for both frequencies of meioses, with mean *π* = 4.01 × 10*^−^*^5^ for *α* = 1 and 4.11 × 10*^−^*^5^ for *α* = 0.01 (Fig 1A). This difference is small, less than 3%, but significant (*p* = 0.00011, one-sided Student’s *t*-test). This similarity between the two cases was expected, because recombination has no effect on the magnitude of neutral variability. The slight difference is due to the fact that, when sexual reproduction is rare, a pair of alleles sampled from the same individual coalesce more slowly than a pair from different individuals, resulting in an increase in *π* [17,18,29].

**Fig 1.**
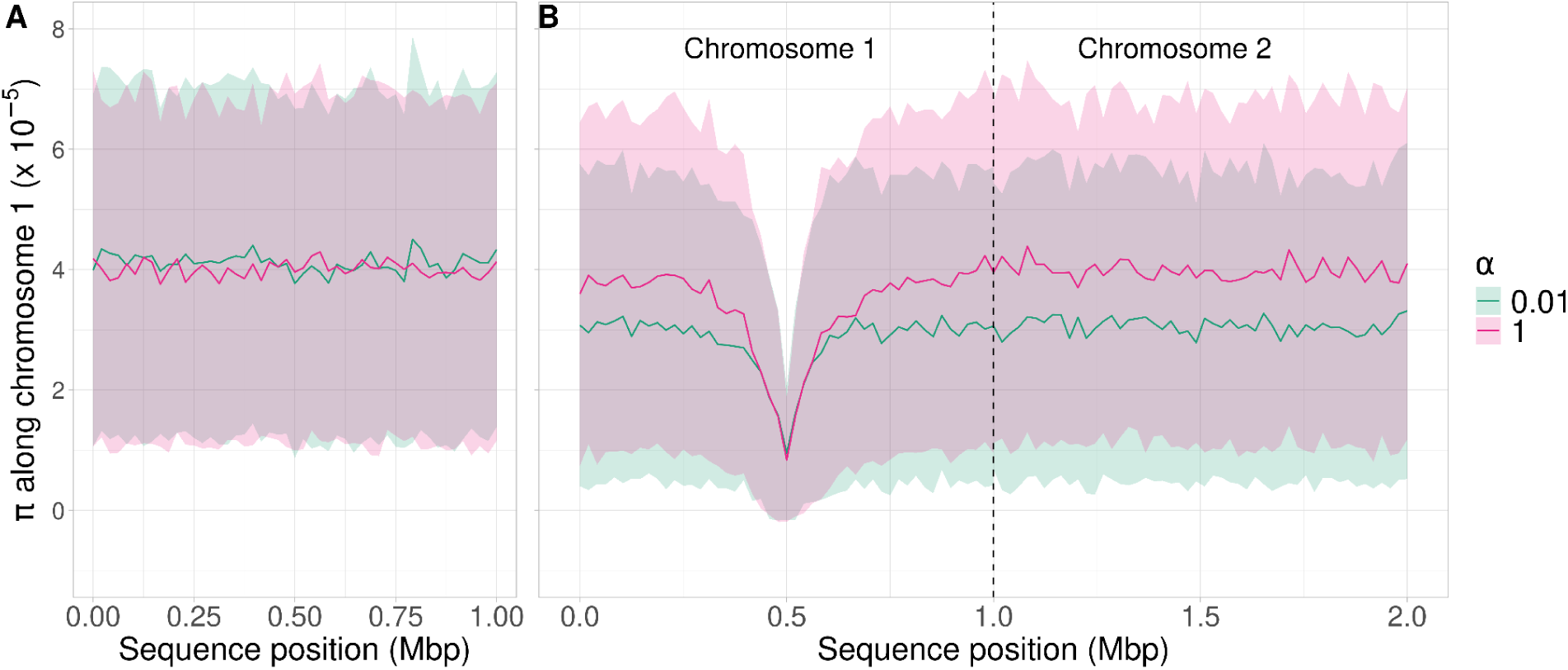
Variation in *π* along a 1 Mb chromosome with *ρ* = 5 × 10*^−^*^8^, and along an independent 1 Mb chromosome. (A) A population of 1,000 individuals that had evolved neutrally for 10,000 generations. (B) A beneficial mutation (*h* = 0.5*, s* = 0.05) at position 0.5 Mb of the first chromosome was introduced into one individual following equilibration under neutral evolution and the population was followed until its fixation. If the mutation was lost before fixation, the simulation was rerun. The second chromosome carried only neutral mutations. In both panels, the pink line is the mean *π* over 500 simulations with *α* = 1, and the green line is the mean *π* with *α* = 0.01, such that the interval between two sexual generations is 1*/α* = 100 generations. *π* is measured over non-overlapping 20 kb windows along the genome. The shaded areas represent standard deviations (see Material and Methods for further details).

When a beneficial mutation was introduced, there was a large loss of genetic diversity around its location, as expected from canonical sweep models [30] (Fig 1B). For both values of *α*, the diversity valley was centered around the sweep, spanning approximately 0.25 Mb on either side of the selected site. Beyond this region, diversity reached a plateau. In addition, *π* ≈ 9 × 10*^−^*^6^ at the location of the selected site, about 22% of the value in the absence of selection (*π* was averaged over 20kb windows, so this value is not exact for the selected site itself). Note that diversity at the selected site is not expected to be zero immediately after fixation, due to recovery of diversity during the sweep (see Equations S1.10 of Appendix S1). The mean numbers of generations taken for the beneficial mutation to reach fixation were similar for the two cases, but the time to fixation, *t,* was significantly higher (*>* 3%) with *α* = 0.01 (*t* = 430 for *α* = 0.01 and *t* = 418 for *α* = 1 with *p* = 0.026, Mann–Whitney *U* test, S1 Fig). The results with a fixed *α* (Fig 1B) did not differ significantly from those with random variation in the interval between meioses (S2 Fig).

The *t* value for *α* = 1 is close to the theoretical value *t_s_* for a semidominant mutation (*h* = 0.5) with homozygous selection coefficient *s* in a sexual population and 2*N_e_s* >> 1, derived by Hermisson and Penning [31]:

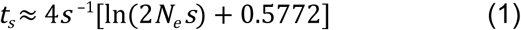

Here *N_e_* is the effective size of the sexual population (*N_e_* = 1,000 in the present case). With *s* = 0.05, *t_s_* ≈ 414.6. In contrast, with *α* = 0.01, *t* is increased to 430 generations, which reflects the fact that fixation requires at least one meiotic event to produce homozygotes for the beneficial allele (see S1 Appendix).

To assess whether the difference in diversity between the two conditions extends further across the genome, we simulated a second chromosome that segregated independently of the first, and which carried only neutral mutations. Diversity on the second chromosome was reduced from 4.01 × 10*^−^*^5^ with *α* = 1 to 3.06 × 10*^−^*^5^ with *α* = 0.01 (*p <* 2.2 × 10*^−^*^16^, one-sided Mann-Whitney *U* test). The diversity plateau seen on the first chromosome was conserved along the second chromosome, so that the fixation of the beneficial allele affected diversity across the entire genome. The effect of selection thus extends across the whole genome when *α* is small (Fig 1B). At first sight, this behavior might seem unexpected, since the net frequency of recombination events per basepair per generation, as measured by *ρ*, is the same for both *α* values. However, the loci on the 2nd chromosome recombine freely with the target of selection at each meiosis, and they are not constrained by *ρ*. Facultative sex thus reduces their effective frequency of recombination with the target of selection to ½*α*, which is the same as the upper limit to the rate of recombination between loci on the first chromosome (see Equation 3 in the Material and Methods).

### How other evolutionary parameters modulate the effect of the frequency of meioses on genome-wide diversity

The extent to which the meiotic frequency *α* and the selection coefficient *s* affect *π* was examined using intermediate values of *α* (0.02 and 0.1) and *s* (0.02 and 0.1). To better reflect natural processes, we incorporated variability in *α* by replacing the fixed interval assumption by sampling the time to the next meiotic event from a normal distribution with mean 1*/α* and a standard deviation of 1/(10*α*) (see Material and Methods). We measured the effect on diversity by dividing *π* by *π*_0_, where *π*_0_ is the baseline neutral diversity, estimated from the mean *π* for chromosome 2 with *α* = 1. The diversity at the sweep position was the same at the end of the simulations for all *α* values and depended only on *s* (Fig 2A-C). This is because the mean times to fixation of the beneficial mutation for a given *s* were mostly similar across the different *α* values; though they sometimes differed significantly, but by less than 3% (S3 Fig).

**Fig 2.**
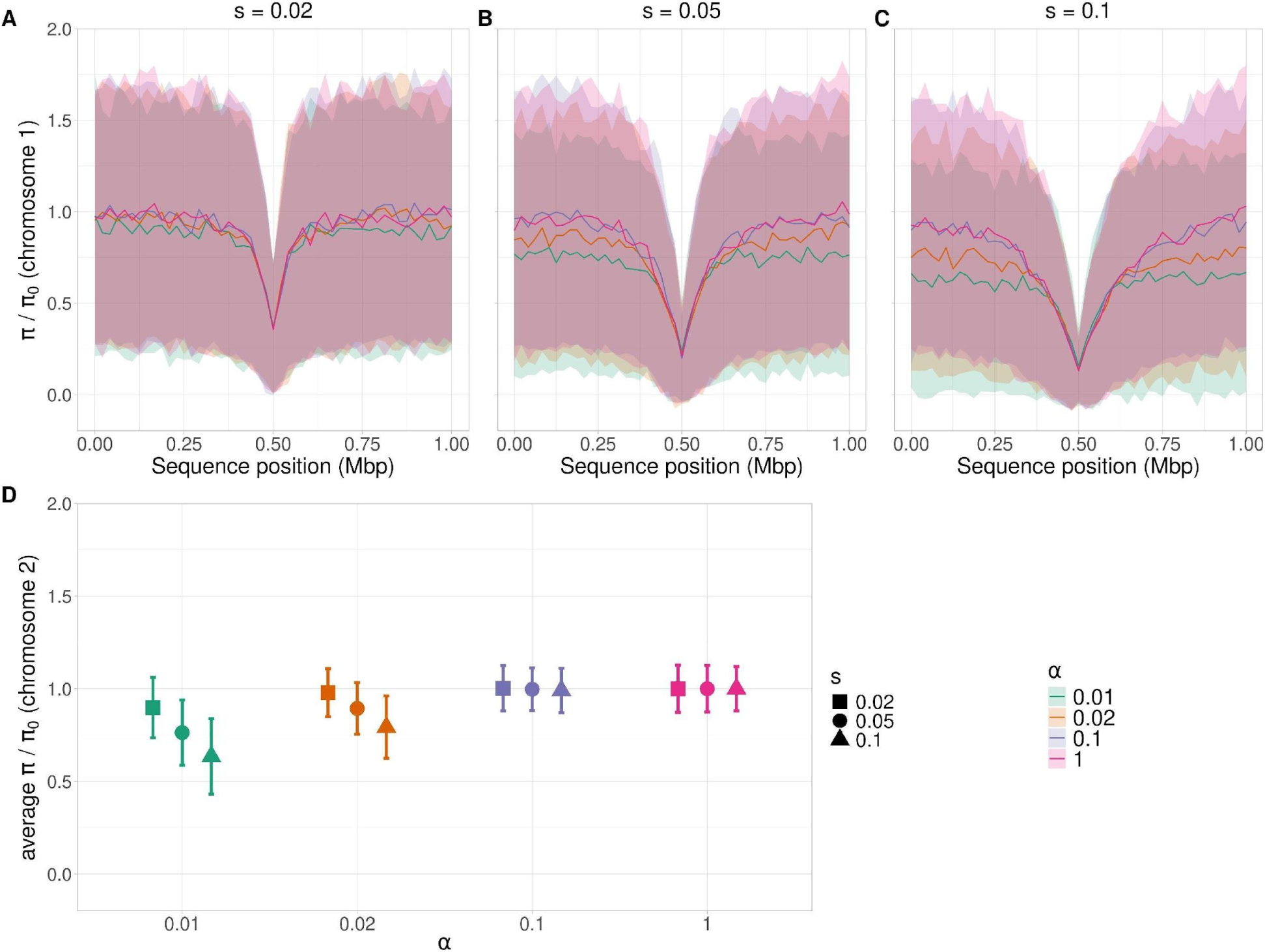
The behavior of diversity relative to the fully neutral value, *π/π*_0_, with different strengths of selection and frequencies of meioses. *π*_0_ is the baseline neutral genetic diversity with *α* = 1 in the absence of selection. Panels A-C show *π/π*_0_ along a 1 Mb chromosome with *ρ* = 5 × 10*^−^*^8^ and a hard sweep at 0.5 Mb; *h* = 0.5 and *s* = 0.02 (A) *s* = 0.05 (B), and *s* = 0.1 (C). Panel D shows mean *π/π*_0_ and its standard error for a second, independent chromosome carrying only neutral mutations. Colors represent *π/π*_0_ with *α* = 0.01 (green), *α* = 0.02 (orange), *α* = 0.1 (blue), and *α* = 1 (pink). The simulation method was the same as in Fig 1, except that the interval between two sexual generations with *α <* 1 was drawn from a normal distribution centered on 1*/α* generations (see Material and Methods for details). *π* is measured over non-overlapping 20 kb windows along the genome. Lines in panels A-C are the means from 500 simulations, and the shaded areas are the standard deviations per site. See Material and Methods for details.

As expected from the established theoretical results on hard sweeps [8,9,19,30,32], the valley of diversity around the target of selection on chromosome 1 becomes narrower as *s* becomes smaller (pink lines in Fig 2A-C), but is relatively unaffected by the value of *α*. In contrast, for *α <* 0.1, *π* on the neutral chromosome decreases with decreasing *α* (*p <* 0.05 for pairwise comparisons, Dunn’s post hoc test) – see Fig 2D. The similarity between the results for *α* = 1 and 0.1 extends to the value of *t,* which suggests that the interval between meiotic events with sufficiently large *α* does not greatly affect the process of fixation of a beneficial allele (S3 Fig). For all *α* values, *t* decreases as *s* increases, as expected from standard theory (*p <* 0.05, Dunn’s post hoc test, S3 Fig). The mean number of recombination events during a sweep for a given *α* increases with larger *t* (see S1 Appendix) allowing *π/π*_0_ to approach one with small *s* and free recombination, even for the lowest *α* value (Fig 2D). In contrast, for large *s* and *α <* 0.1, recombination events are too rare for diversity to fully recover.

In order to interpret these findings, we compared the simulation results presented in Fig 2 to the theoretical predictions of the effects of a sweep in a randomly mating population, given by Equations 14 of [32]. For this purpose, we used the product of *α* and the recombination rate *r* per meiosis between a neutral locus and the target of selection (where *r* is given by Equation 3 in the Material and Methods) in the relevant formulae. This provides a measure of the net rate of recombination per generation between the two loci, and should not be confused with *ρ,* the net frequency of recombination per basepair per generation. For *α* < 1, the standard approximate equation for change in allele frequency at an autosomal locus, Δ*q* ≈ *sq*(1 – *q*)[*h* + (1 – 2*h*)*q*], needs to be modified to Δ*q* ≈ *sq*(1 – *q*)[*αh* + 2(1 – *α*)*h*+ *α*(1 – 2*h*)*q*], to take into account the fact that asexual diploids behave like haploids with selection coefficient *2hs* (see section 2 of S1 Appendix). This procedure is likely to only be an adequate approximation when selection is weak and *α* is sufficiently large that the population can be treated as though it is partially sexual and partially asexual at all times, which is far from being true with small *α*.

As expected from previous results [32], the model performs well for all three selection coefficients (*s*) when *α* = 1 (Fig 3 A-C), as this represents the “canonical” case for selective sweeps. The use of *rα* as a straightforward measure of an effective recombination rate leads to a reasonable approximation, especially for *s* = 0.02. However, the fit deteriorates as *α* decreases and s increases, and the model tends to systematically underestimate diversity, highlighting its limitations in regimes of infrequent meiosis. In the parameter space considered here (*h* = 0.5 and *α* ≥ 0.01), the model from [32] captures the qualitative trend of diversity reduction, but a more refined theoretical framework is needed to accurately predict the magnitude of diversity loss with low *α* (see the next section). This finding suggests that while incorporating *α* into the recombination rate is a useful first step, standard sweep theory alone does not fully capture the dynamics of facultative sex. The plots for different *α* values, but the same *s,* in Fig 2 converge when the distance becomes small. The convergence reflects the fact that the recombination rate obtained from equating the product of *ρ* and distance to *d* in Equation 3 is the same for all *α* when *d* is small, because the recombination rate is then close to the map distance.

**Fig 3.**
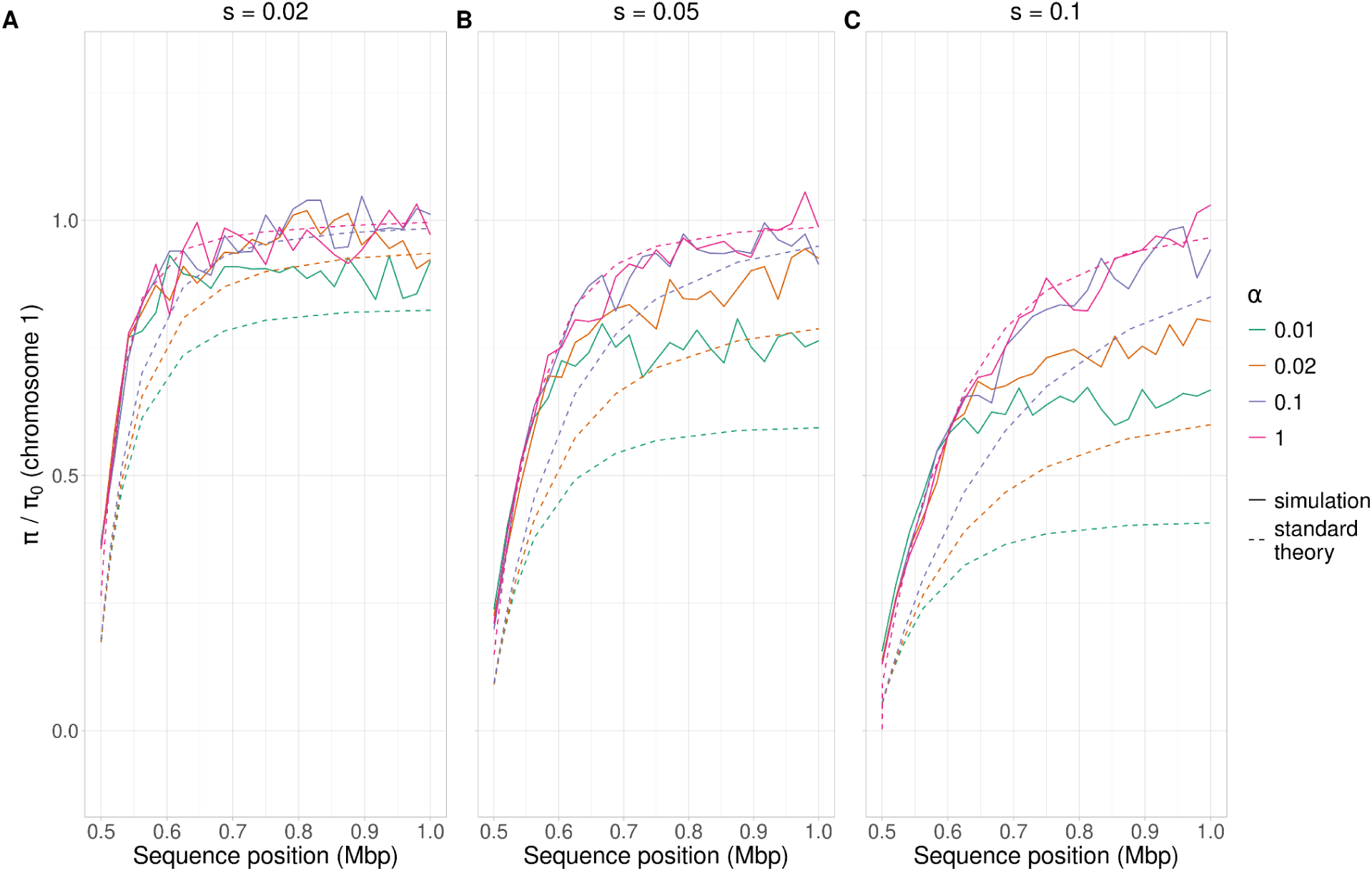
Comparison of the properties of *π/π*_0_ between theory and simulations. *π/π*_0_ along a 0.5 Mb chromosome (see Methods) with a hard selective sweep at 0.5 Mb; *h =* 0.5 and *ρ* = 5 × 10*^−^*^8^ (A) *s* = 0.02. (B) *s* = 0.05 (C) *s* = 0.1. “Standard theory” corresponds to Equations 14 from [32], with the recombination frequency from Equation 3 rescaled as *rα* and using the selection equation described in the text (dashed lines). Simulated data are the averages from 500 simulations (solid lines). Colors represent diversity relative to the fully neutral case, with *α* = 0.01 (green), *α* = 0.02 (orange), *α* = 0.1 (blue), and *α* = 1 (pink). *π* is measured over non-overlapping 20 kb windows along the genome.

Additionally, we investigated how the recombination rate per base (*ρ_α_*) and the dominance coefficient (*h*) influence genetic diversity during a selective sweep. *ρ* rather than *ρ_α_*primarily dictates the width of the genetic diversity valley around the sweep (S4 and S5 Figs, sections 1 and 2 of S3 Appendix). The dominance coefficient *h* has a minimal effect on genetic diversity with *α* = 1, but its influence becomes more pronounced at lower *α*, with higher *h* (a stronger heterozygous effect of a beneficial mutation) causing increased diversity, as is also seen for a sexual population (Fig 1 of ref [32]). We also compared the simulation results with the theoretical approximation for large *α* (as shown in Fig 3) for different values of *h* (S7 Fig). While the fit remains good for *α* = 1, it deteriorates for lower *α* values and becomes increasingly inaccurate as *h* deviates from 0.5. The reasons for these discrepancies are not obvious but must reflect the crude nature of the approximation used here. A detailed analysis of the effects of both evolutionary parameters *h and ρ_α_* is given in sections 2 and 3 of S3 Appendix.

### Recovery of diversity during a sweep with a single meiotic event

The standard sweep theory does not accurately predict the genetic diversity (*π*) for low *α* values, as shown in Figs. 3 and 4. In these cases, the standard sweep theory using *rα* as an effective recombination rate strongly underestimates *π*. The analysis of the case of *α* << 1 in the S1 Appendix shows that there is a “phase shift” between the behavior of the system with large and small *α* values, which can be understood as follows. For a neutral site with a recombination frequency with the selected locus of *r* per meiosis, a single meiotic event involving the heterozygous carriers of the beneficial allele has a probability of 2*r*(1-*r*) of producing homozygotes for the beneficial allele that are non-identical by descent at the neutral site (section 1 of S1 Appendix). Since these homozygotes ultimately replace the other genotypes if there are no further meiotic events, a scenario with at most one meiosis during the initial sweep of heterozygotes for the beneficial allele would give *π / π*_0_ = 0.5 when *r* takes its maximal value of ½, all other things being equal.

**Fig 4.**
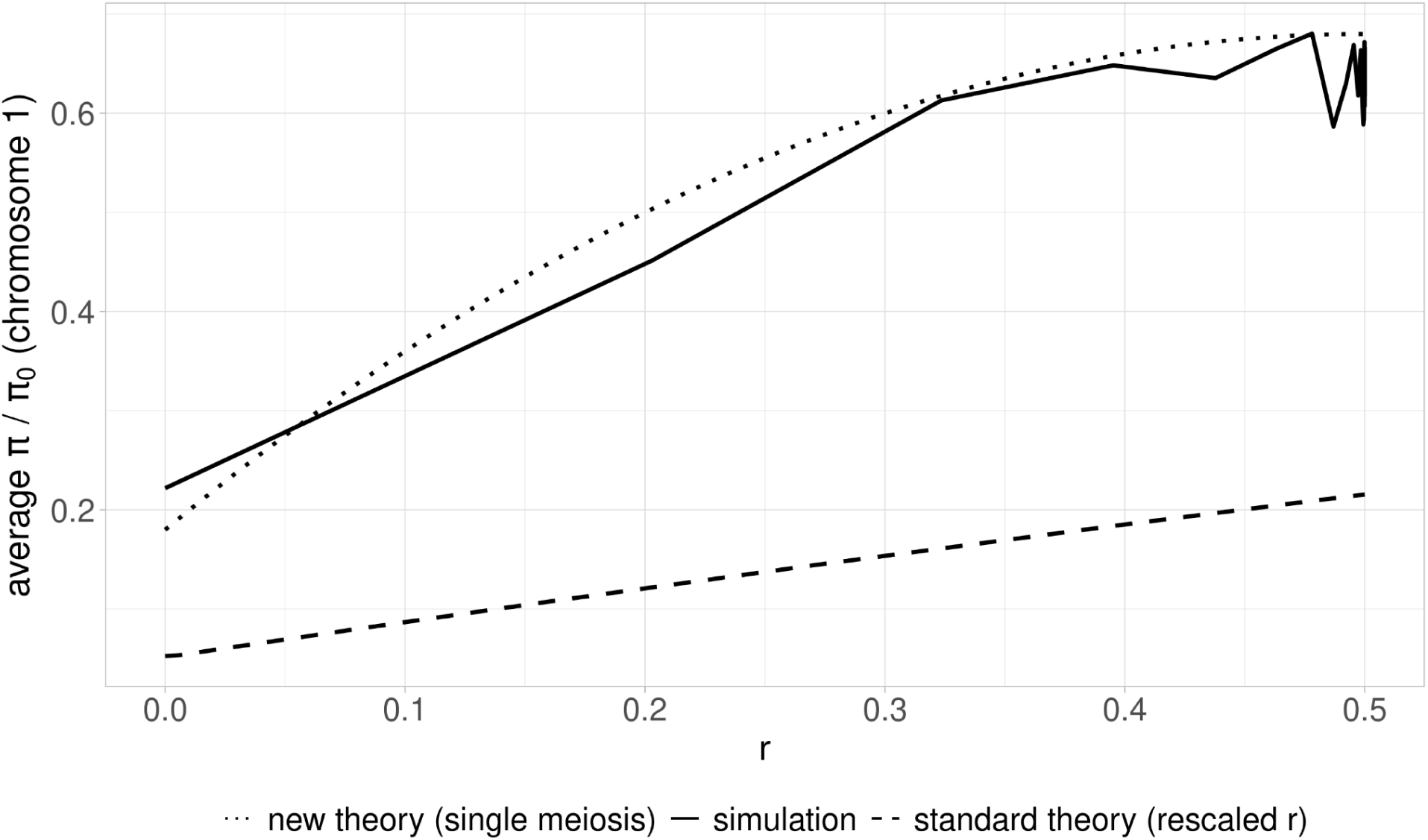
The relation between *π / π*_0_ and *r,* the recombination frequency per meiosis between the neutral and selected loci, where *r* is given by Equation 2 in Material and Methods). The lines represent the predicted and simulated values of *π / π*_0_ for small *α* (0.004). The theoretical values are from Equation S1.10 (dots) or from Equations 14 of [32] with *r α* as the effective recombination rate (dashes). The solid line shows the mean values of *π / π*_0_ over 500 simulations. A beneficial mutation (*h* = 0.5*, s* = 0.1) was introduced into one individual at one end of a 0.5Mb chromosome with a recombination rate per basepair of *ρ* = 5 × 10*^−^*^8^, and the population allowed to evolve until fixation of the beneficial mutation, as in previous figures. The time interval between two sexual generations was drawn from a normal distribution centered on 1*/α* generations. See Material and Methods for details.

However, diversity will increase above this value, due to the accumulation of new mutations at the neutral site (section 2 of S1 Appendix), an effect that is also included in Equations 14 of [32]. This increase is amplified by the fact that, once the meiotic event has occurred, there is a second time interval during which homozygotes for the beneficial allele increase in frequency and become fixed (see Equations S1.5 and S1.8 of S1 Appendix). When multiple meiotic events occur during the sweep, the final *π / π*₀ ratio will be much higher than predicted by the model based on a single event (Equations S1.10), and the numbers of such events can be approximated by the product of *r, α* and the sweep duration, as in the standard sweep model. An approximate analysis of the case of two meiotic events per sweep is given in S2 Appendix; there is, however, only a relatively narrow range of parameter space to which this model applies.

Fig 4 compares simulation results with small *α* and large *s* values with the predictions of Equations S1.10 of the S1 Appendix and those of standard sweep theory. With *s* = 0.1, *h* = 0.5, and *ρ* = 5 × 10*^−^*^8^, the expected time for the initial sweep of heterozygotes for the beneficial allele to go to completion is *t_s_*_1_ = 207.2 generations (Equation 1). With *α* = 0.004, the expected number of meioses per sweep is given by *α t_s_*_1_= 0.829. Since the requirement of at most one meiotic event during the initial sweep is satisfied, the analytical predictions align well with the simulation results, in contrast to the predictions of standard sweep theory.

## Discussion

This study is the first to analyze the effects of a hard selective sweep in a diploid species that reproduces by facultative sex, with long intervals between episodes of sexual reproduction that occur simultaneously in all individuals in the population. We acknowledge that the parameter space explored here is restricted, focusing on scenarios with two chromosomes, a single selective sweep, and a limited range of meiotic frequencies (*α*). Contrary to what is expected from standard models, the effect of a sweep on diversity for a given strength of selection is not simply determined by the net frequency of recombination between a neutral site and the target of selection (*i.e*, by the product of the recombination rate per meiosis and the frequency of sexual generations, *α*), except when the frequency of meioses is sufficiently high that several sexual generations occur during a sweep. Otherwise, the expected diversity at a given neutral site is controlled jointly by the probability that a single smeiotic event generates homozygotes for the beneficial allele that are not identical by descent at the neutral site and by the time available for diversity to accumulate during the sweep (see Fig 4, the S1 Appendix, and the S2 Appendix). Under these conditions, a selective sweep can influence not only nearby loci, but also the entire genome, if the number of meioses per sweep is sufficiently small.

While our model deliberately simplifies certain aspects of natural systems, by focusing on hard sweeps in an outbreeding diploid organism with only two chromosomes, many of the qualitative patterns we observe are likely to generalize. For example, the potential for genome-wide reductions in diversity due to rare recombination during selective sweeps is expected to apply in any facultatively sexual organism with a low meiotic frequency (see sections 4 and 5 of the S1 Appendix and section 4 of S3 Appendix). However, other effects, such as the shape of the *π/π*₀ curve under specific dominance values, or the precise point at which standard sweep theory breaks down, depend on the chosen parameters, including population size, chromosome architecture, and selection strength. We emphasize that the analytic approximations derived here are only accurate for restricted regions of parameter space and should be interpreted accordingly. Thus, while we believe our results highlight robust and underappreciated effects of rare sex on genome-wide diversity, they should be seen as a first step toward understanding more complex and realistic scenarios.

The rescaled standard model, where the recombination rate between the selected and neutral locus (*r*) is multiplied by *α*, can be most reliably applied when *h* = 0.5 and *α* ≥ 0.1, provided that *αt_s_*>> 1, where *t_s1_* is the duration of the initial sweep of heterozygotes for the beneficial mutation given by Equation S1.3b, but it is less accurate when *h* deviates from 0.5. The use of *rα* in the sweep equations is reminiscent of the use of *r* (1 – *F*) as an approximate effective recombination rate with an inbreeding coefficient of *F* in standard sweep theory [32]. In contrast, Equations (S1.10) seem to give accurate analytical predictions for situations in which at most one meiosis occurs during a sweep. Interestingly, the major role of 2*r*(1 – *r*) in Equations S1.10 violates the widely used principle that properties such as *π / π*_0_ should be determined by the products of *N_e_* and deterministic parameters such as *r*, rather than their absolute values [8]. This reflects the fact the process controlling diversity when there is at most one meiosis per sweep is largely deterministic rather than stochastic.

These results raise the broader question of how the values of parameters such as *α*, *ρ* and *r* used in the simulations with a population size of *N_s_* = 1,000 can be related to their values in a natural population with a much greater size, *N_n_*. When *αt_s_*_1_ is >> 1 and standard sweep theory is applicable, at least as an approximation, the scaled parameters 2*N_e_s* and 2*N_e_αr* determine the effect of a sweep on *π / π*_0_. When this is the case, the simulation value of *α* would also apply to the natural population, whereas *s* and *r* should be rescaled by multiplication by *N_s_*/*N_n_* to capture their natural population values (fixation times are correspondingly multiplied by *N_n_*/*N_s_*). In contrast, when *αt_s_*_1_ is ≤ 1, standard sweep theory no longer applies, due to the rarity of recombination events. In this case, Equation S1.10b implies that *r* should not be rescaled, whereas *α* should be multiplied by *N_s_*/*N_n_*. This leads to the somewhat unsatisfactory conclusion that the relations between the simulation results and the real world are dependent on the value of *αt_s_*_1_ or, equivalently, the product of the scaled parameters 2*N_e_α* and *T_s_*_1_ = *t_s_*_1_ /2*N_e_.* Since *t_s_*_1_ is largely determined by the reciprocal of *hs*, the effects on diversity of strongly selected beneficial mutations are likely to be largely controlled jointly by *r* and 1/(2*N_e_α)* (Equation S10b), whereas the effects of weakly selected mutations are more likely to be controlled by 2*N_e_s* and 2*N_e_αr*.

Further work is required to extend our theoretical understanding to situations where *α t_s_*_1_ is not too far from 1, where the transition to diversity that is largely controlled by the recombination rate *r* and 1/(2*N_e_α*) takes place. In addition, the effects of recurrent sweeps need to be studied, together with population genetics statistics such as levels of linkage disequilibrium, the extent of distortions of the shapes of gene genealogies by the effects of selection at linked sites, and the extent of Hill-Robertson interference among the selected sites themselves. In addition, the model could be extended to incorporate biological complexities such as multiple simultaneous sweeps, spatial population structure, and more realistic life cycles. For example, some facultatively sexual species exhibit parthenogenesis or haplo-diploid cycles with prolonged haploid clonal phases, as seen in algae like *Chlamydomonas* [33]. Our model avoids these complexities by focusing solely on a diploid population reproducing either clonally or sexually via outcrossing, which allows us to isolate the core effects of facultative sex on diversity and sweeps. The model can readily be extended to a diploid clonal phase with inbreeding (section 4 of the S1 Appendix) and a haploid clonal phase with a transient diploid zygote (section 5 of the S1 Appendix). We ran simulations with various selfing rates (S11 Fig and section 4 of S3 Appendix). These features result in larger sweep effects than those described here, extending over larger regions of the genome, as would be expected from the reduced effectiveness of recombination with selfing.

It is of interest to compare the life cycle modeled here to that of multicellular organisms. Under neutrality, both life cycles involve multiple mitotic divisions before sex and may thus generate similar patterns of diversity. However, in the presence of selection, important differences arise: selection acts primarily on individual cells in unicellular or simple multicellular organisms, whereas it acts primarily at the organism level in species with a differentiated germline.

The motivation for emphasizing *Saccharomyces cerevisiae*, as is done here, stems from its composite life cycle and its status as a model organism. As a facultatively sexual species, *S. cerevisiae* and its relatives alternate between clonal reproduction and sexual reproduction, making it ideal for exploring the impact of meiotic frequency on patterns of genetic diversity. Although this study focuses exclusively on the effects of outcrossing events, we acknowledge that other modes of sexual reproduction such as haploselfing (the intra-tetrad mating of the products of meiosis) also play a role in shaping genetic diversity (see section 4 of S3 Appendix and S11 Fig). Many evolutionary parameters for this species, including mutation rates, are well established through laboratory experiments [25,34,35]. Additionally, the large effective population sizes of *S. cerevisiae* and its relatives ensure that natural selection should operate efficiently, providing a robust framework for studying the genome-wide effects of selective sweeps. With the increasing availability of genomic data, addressing questions about the persistence of rare recombination events in yeasts [12,36] could become another avenue for exploration. Understanding whether these events contribute to maintaining genetic diversity or if they are simply evolutionary byproducts could help explain the evolutionary advantage of maintaining recombination. Moreover, our findings are relevant to other facultatively sexual species that alternate between clonal and sexual reproduction, such as *Daphnia*, *C. elegans*, and bdelloid rotifers, or to species such as *S. paradoxus*, in which sexual reproduction is extremely rare (e.g., *α* ≈ 10*^−^*^4^ in some lineages [14,15]). Strikingly, even within a single species like *S. cerevisiae*, estimates of the frequency of sex can vary by more than 100-fold across natural populations, as shown for Taiwanese lineages by Lee et al. [12]. These systems provide valuable opportunities to test whether rare outcrossing events can have genome-wide effects due to limited recombination during sweeps.

Studies on wild *S. paradoxus* have provided estimates of rates of clonal reproduction as high as 0.9999 (*α* = 0.0001) [14,15], but this assumes that neutral diversity at synonymous sites is not influenced by selection. Here, we provide evidence that selection can alter global genetic diversity, highlighting the importance of considering selective effects in similar analyses. A limitation is that our study focuses on hard sweeps and does not take into account the fact that most mutations in *S. cerevisiae* are probably nearly neutral or deleterious [37]. It would be desirable for future work to explore the effects of negative selection on genome-wide genetic diversity (*π*), including the effects of background selection (BGS) [38] and associative overdominance (AOD) [39,40]. These processes, which either decrease or increase genetic diversity respectively, are influenced by recombination rates and meiotic frequency, with AOD becoming more prominent under low recombination conditions. Understanding the transition from BGS to AOD, and its dependence on *α* and the recombination rate per meiosis remains an area for further investigation.

This study also provides a new perspective on Lewontin’s paradox—the observation that genetic diversity across species of eukaryotes does not increase as fast with census population size as expected from neutral theory [22]. While previous research has attributed this discrepancy to factors like historical changes in effective population size [23,24], our findings suggest that differences in breeding systems, particularly meiotic frequency, could also play a significant role. Species with a low meiotic frequency, even those with large census sizes, may exhibit reduced genetic diversity compared with species that reproduce fully sexually. However, we acknowledge that Lewontin’s paradox remains unresolved, and that species with similar reproductive strategies and recombination frequencies can show wide variation in diversity levels [23]. This highlights the need to estimate meiotic frequency and analyze its relationship with census size and *N_e_* across species to better understand the dynamics of genetic diversity. By focusing research efforts on meiotic frequency, we can further elucidate the factors driving patterns of genetic variation across diverse populations. Infrequent sex creates a regime where genome-wide linkage allows a single selective event to reshape diversity patterns far beyond the selected locus. This implies that seemingly neutral genome-wide diversity can bear the signature of rare selective events, with important implications for interpreting genetic variation in facultative sexual organisms. As discussed above, the mean number of recombination events during a sweep increases with longer sweep durations, allowing *π*/*π*_0_ to approach one when *hs* is small, even for the lowest *α* value (Fig 2D). In contrast, for large *hs* and α < 0.1, recombination events are too rare to restore diversity to fully recover. By explicitly incorporating variation in meiotic frequency into models of the effect of selection at linked sites, we may improve our understanding of how reproductive strategies contribute to shaping genetic diversity across taxa.

## Material and Methods

### Modeling reproduction and meiotic recombination

Two new parameters are introduced compared to canonical population genetic models: the recombination rate per site per sexual generation, *ρ_α_*, and the frequency of sexual reproduction (*i.e*, meiosis), *α*. We assume an absence of mitotic recombination, as it is four orders of magnitude less frequent than meiotic recombination in yeast [34,41]. In most simulations, we assumed a constant average recombination rate per nucleotide site per generation, weighted by the frequency of meiotic events:

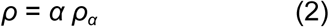

The motivation for this procedure is to examine the extent to which the size of the reduction in diversity caused by a sweep is determined by the ratio of the recombination rate per meiosis, modulated by the frequency of meioses to the strength of selection on the beneficial allele, as expected from classical sweep theory (where *α* = 1) [30,32]. When *α <* 1, reproduction occurs both sexually and asexually; at each generation, all diploid individuals reproduce in the same way. For example, when *α* = 0.01, meiosis and recombination occur on average once every 100 mitotic divisions. We used values of *α* = [1, 0.1, 0.02, 0.01] in most simulations.

We modeled the occurrence of sexual generations when *α <* 1 using two approaches. Initially, sexual generations were set to occur at fixed intervals of 1*/α* generations. Later, this assumption was relaxed to allow variability in *α*. In that case, we determined the next sexual generation by sampling from a normal distribution centered on *x* (where *x* = 1*/α*) with a standard deviation *σ* = *x/*10. This approach enabled the exploration of both regular and stochastic patterns of sexual reproduction.

### Simulation methods

We performed forward-in-time, individual-based simulations using SLiM [4]. In a sexual generation, two diploid parents were sampled from the population with a probability proportional to their fitness and allowed to mate to produce a single diploid offspring; this procedure was repeated until *N* individuals were formed. In an asexual generation (clonal reproduction), *N* offspring were produced by sampling a single parent, drawn from the population in proportion to its fitness. This is a Wright-Fisher sampling scheme, where the effective size of the population is *N* for a sexual generation. In contrast, for asexual generations, the effective population size is approximately 0.5*N*, because both chromosome copies are inherited from a single parent without recombination, which increases the rate of coalescence and reduces the level of genetic variation to values corresponding to a haploid population of size 0.5*N* [8].

We chose the values of the mutation rate [26] and recombination rate per basepair [34] to mirror *S. cerevisiae* evolutionary dynamics, using the principle that the products of effective population size (*N_e_*) and deterministic parameters control the dynamics of evolution if time is measured in units of 2*N_e_* generations [8,42]. *N_e_*was estimated by plugging the *π* value from [28] and mutation rate from [26] into the formula: *N_e_ = π/4µ*. We thus modeled a plausible natural population effective size of 10^6^ as far as diversity is concerned. Caution is needed when applying these results to a natural population of this size, as the recombination rates per meiosis for closely linked sites in the simulations are probably much lower than those that are appropriate for modeling a natural population, except when *α* is very small, given that the mean rate of recombination per Mb in yeast is approximately 0.005 per Mb (0.004 in [43] and 0.0061 in [34]). We simulated two 1Mb independent chromosomes for 1,000 diploid individuals using the reproduction scheme described above. The second chromosome carries only neutral mutations.

In the absence of selection, simulations were run for 10,000 generations. When natural selection was included, a site under strong positive selection was introduced into the middle of the first chromosome of one individual at generation 2,000. Selection acts at each cell division (both mitoses and meioses). The selection and dominance coefficients, *s* and *h*, determine the fitnesses of the mutant genotypes relative to wild-type as 1 + *hs* for heterozygotes and 1 + *s* for homozygotes. Unless specified otherwise, the beneficial mutation was semidominant, with *h* = 0.5 and *s* = 0.05 (2*Nhs* = 50). Simulations were stopped when the beneficial mutation reached fixation, when all individuals in the population were mutant homozygotes. In the rare cases where the mutation failed to fix before 10,000 generations, the simulations were run again from the beginning until fixation occurred. 500 replicates for each parameter set were run.

To estimate diversity at the neutral sites, we retrieved tree sequences and the generation when fixation occurred from the simulation output [44]. Following the most recent recommendations of SLiM [4], we used tskit (https://github.com/tskit-dev/tskit), pyslim (https://github.com/tskit-dev/pyslim), and msprime [45] Python (version 3.8.13) packages to get fully coalesced population trees (the recapitulation step) and to remove extinct lineages (the simplification step). We then added neutral mutations to the resulting tree (the overlay step) with a neutral mutation rate per site per generation of *µ* = 1 × 10*^−^*^8^.

### Summary statistics

For each simulation, we sampled 100 individuals from the population at the time of fixation of the beneficial allele, and computed the average nucleotide site diversity at neutral sites for the sampled individuals [46] using tskit (https://github.com/tskit-dev/tskit) Python package. We used the site mode to estimate the mean pairwise genetic diversity for 50 windows along the first chromosome (20kb non-overlapping windows) and for the whole second chromosome. We computed the mean and standard deviation of *π* for each window between the 500 replicates. We quantified the reduction in genetic diversity for a given parameter set by dividing *π* by *π*_0_, where *π*_0_ is the baseline neutral diversity in the absence of sweeps, estimated from the mean *π* for chromosome 2 when *α* = 1.

### Mathematical modeling

An analytical model with an arbitrary value of *α* is hard to construct, due to the complex dynamics when there is more than one meiosis during a sweep. However, if the product of *α* and the duration of a sweep is sufficiently low that at most a single meiotic event occurs during a sweep, a relatively simple formula can be obtained for the expected neutral diversity at a site which recombines with the selected site with frequency *r* per meiosis – see Equations (S1.10) of S1 Appendix. The simulation method assumed a complete lack of crossover interference, so that *r* in the simulations is related to the map distance *d* between the neutral and selected sites by Haldane’s mapping function [47]:

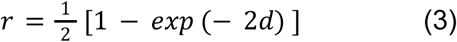

where *d* is equal to the distance from the sweep in base pairs multiplied by the recombination rate per site per sexual generation, given by *ρ_α_* = *ρ*/*α*.

### Statistical tests

Differences in mean *π* values on the second (neutral) chromosome between two different *α* values were tested by first checking that they follow a normal distribution (Shapiro test), either using a Student’s *t*-test (if Gaussian) or Mann-Whitney *U* test (if not). We compared the average time to fixation of hard selective sweeps between two conditions by the same method. When comparing more than two cases at once, we performed Kruskal-Wallis tests followed by a Dunn’s post hoc test [48] (these cases were never all Gaussian). All these statistics were analyzed using R (version 4.3.2) [49].

### Figs

Plots were made using the following packages from R (version 4.3.2) [49]: cowplot [50], ggplot2 [51], readr [52], tibble [53], dplyr [54] and RColorBrewer [55]. The colors should be color-blind-friendly.

### Cluster computations

Simulations were run on a SLURM-based (https://github.com/SchedMD/slurm) HPC cluster at IFB (https://www.france-bioinformatique.fr/cluster-ifb-core/) within the software environment management system conda (https://github.com/conda) for reproducibility. Scripts are available here: https://doi.org/10.5281/zenodo.15657189.

### Data Availability

All scripts to perform the simulations and make the figures, the results as well as a conda environment are available here: https://doi.org/10.5281/zenodo.15657189.

## Acknowledgments

The simulations were performed on the Core Cluster of the Institut Français de Bioinformatique (IFB) (ANR-11-INBS-0013). We would like to thank George Marchment for correcting the English as well as Florian Massip, Laurent Excoffier, Guillaume Achaz, Gilles Fischer, Alexander Palazzo and Nina Vittorelli for helpful discussions about the project.

## Supporting Information caption

**S1 Appendix. The effect of a selective sweep with only one meiosis per sweep**

**S2 Appendix. Model of a sweep with two meiotic events**

**S3 Appendix. Additional analysis of parameters**

**S1 Supplementary figures**

