## Supplementary material for "Beyond Recombination: Exploring the Impact of Meiotic Frequency on Genome-wide Genetic Diversity": S1 Appendix

### The effect of a selective sweep with only one meiosis per sweep

**1. Model of a sweep of a heterozygote for a beneficial mutation when meiosis is rare …….. *p. 2***

**2. Recovery of diversity during a sweep: no meiosis during the sweep of $A_2 A_1$ …………….. *p. 4***

**3. Recovery of diversity during a sweep: a single meiosis during the sweep of $A_2 A_1$ ……… *p. 7***

**4. Model of a sweep of a heterozygote for a beneficial mutation when meiosis is rare: inclusion of inbreeding ………………………………………………………………………………………… *p. 9***

**5. Model of a sweep in an organism where the asexual phase is haploid …………………….. *p. 11***

**1. Model of a sweep of a heterozygote for a beneficial mutation when meiosis is rare**

The model assumes that the frequency of meiotic events is low, so that $\alpha << 1$; in the absence of meiosis, diploid individuals reproduce clonally. Assume that a beneficial mutation $A_2$ at a selected locus with dominance coefficient h ($0 < h < 1$) and homozygous selective advantage $s > 0$ arises when the population is reproducing clonally, and is initially present as a heterozygote with the ancestral allele $A_1$. Assume also that there is a neutral locus B, which is linked to the A locus with recombination frequency $r$. The state of the sweeping genotype is denoted by $A_2B_2/A_1B_1$, where $B_1$ and $B_2$ are not necessarily different in state, but are initially non-identical by descent (non-i.b.d.). In the absence of a meiotic event, the frequency of $A_2B_2/A_1B_1$ relative to the $A_1A_1$ ancestor obeys the standard equation for selection on a haploid system (e.g. [1] p.89), but with selection coefficient $hs$.

If meioses occur with a fixed interval $\alpha^{-1}$ between successive events, where $\alpha$ is the frequency of meiotic events per generation, we can assume that a sweep occurs at a random time in relation to a meiotic event. Then the p.d.f. for the time $t_m$ to the first meiosis after the start of the sweep is a uniform distribution over the interval $(0, \alpha^{-1})$, such that the probability of an event in the infinitesimal time interval $dt_m$ is $\alpha\, dt_m$. Let the time taken for $A_2B_2/A_1B_1$ to go to fixation be $t_{s1}$. If $t_{s1} < \alpha^{-1}$, the probability that no meiosis occurs during a sweep is given by:

$$P\left(t_m \geq t_{s1}\right) = \alpha \int_{t_{s1}}^{\alpha^{-1}} dt_m = 1 - \alpha t_{s1} \qquad \text{(S1.1)}$$

The expected time to a meiotic event after the beginning of a sweep is given by:

$$\bar{t}_m = \frac{1}{2}\alpha^{-1} \qquad \text{(S1.2)}$$

First, consider the case when no meiotic event occurs during a sweep. If we condition on $A_2B_2$/ $A_1B_1$ having gone to fixation, the probability of i.b.d. at the B locus is ½. For the population to become fixed for $A_2/A_2$, however, at least one meiotic event is needed, in order to allow segregation. A meiotic event involving all individuals in the population allows matings of the type $A_2B_2/A_1B_1$ x $A_2B_2/A_1B_1$ to occur, the equivalent of a standard F2 cross.

If we then condition on random $A_2/A_2$ genotypes becoming fixed, as happens with high probability if $h < 1$ (so that $A_2/A_2$ outcompetes $A_2/A_1$), it is easy to see that the frequencies of $A_2B_2/A_2B_2$, $A_2B_2/A_2B_1$ and $A_2B_1/A_2B_1$ are $(1 - r)2$, $2r(1 - r)$ and $r2$, respectively. The frequency of $B_1$ and $B_2$ among these genotypes are $r$ and $1 - r$, respectively, as might be expected intuitively, so that the probability of non-i.b.d. at B is $2r(1–r)$, assuming that the population size is sufficiently large that the effects of drift can be neglected during the sweep. This probability obviously ranges from 0 ($r = 0$) to ½ ($r = ½$).

Now assume next that a single meiotic event occurs before $A_2B_2/A_1B_1$ has become fixed in the population. In this case, two types of mating are possible: an $F_2$ of the same type as before, and a backcross between $A_2B_2/A_1B_1$ and the $A_1A_1$ population. The backcross will not, however, produce any $A_2A_2$ genotypes, and need not be considered if we condition on $A_2A_2$ having gone to fixation. Thus, this situation produces the same final result as the previous one, as far as the probability of identity by descent at the B locus is concerned.

### 2. Recovery of diversity during a sweep: no meiosis during the sweep of $A_2 A_1$

Identity by descent is not equivalent to lack of diversity at the B locus, because new mutations can accumulate during the process of fixation of $A_2$, so that i.b.d. alleles can become different in state. To investigate this process, we first consider the case of no meiosis during the sweep of $A_2B_2/A_1B_1$ to fixation. Assume that the frequency of meiosis in the pre-sweep population is sufficiently high that the coalescence time of a pair of neutral alleles is $2N_e$, where $N_e$ is the long term effective size of the ancestral population (note that $N_e \approx N$ under the Wright-Fisher model used in the simulations, provided that $\alpha >> 1/N$). During the time that the population reproduces clonally, however, the variance effective population size with respect to the frequencies of $A_2A_1$ and $A_1A_1$ is $N/2$, because the variance in in the change per generation in the frequencies of two alternative genotypes with frequencies $x$ and $(1 - x)$ is $x(1 - x)/N$.

During the process of fixation of $A_2B_2/A_1B_1$, the two B alleles will accumulate mutations independently of each, so that they will come to diverge by an amount relative to the ancestral pairwise diversity ($\pi_0$) that is equal to $T_{s1}$ where $T_{s1}$ is the time to fixation of $A_2B_2/A_1B_1$ scaled by $2N_e$, i.e, $T_{s1} = t_{s1}/(2N_e)$, where $t_{s1}$ is the absolute time to fixation (uppercase letters for times indicate scaling by $2N_e \approx 2N$). This measures the expected increase in diversity relative to the purely neutral expectation.

As pointed out in section 1, provided that $0 < h < 1$, the sweep of $A_2B_2/A_1B_1$ to fixation corresponds to the spread of an allele with selective advantage $hs$ in a haploid population. With sufficiently weak selection ($hs << 1$), this is equivalent to the spread of a semidominant allele with selective advantage $2hs$ in a diploid, randomly mating population. Using the fact that during this period the variance effective population size is equal to $N/2$, we can apply Equation (A.17) of Hermisson and Pennings (2005) [2] for the semidominant case to write:

$$T_{s1} \approx 2\gamma^{-1}[\ln(\gamma) + 0.5772] \qquad \text{(S1.3a)}$$

where $\gamma = 2Nhs$.

The unscaled version of this expression is:

$$t_{s1} \approx (hs)^{-1}[\ln\ (\gamma)\ +\ 0.5772] \qquad \text{(S1.3b)}$$

The increase in diversity during the sweep of $A_2/A_2$ after the fixation of $A_2/A_1$ also needs to be considered. This can only occur after a post-sweep meiotic event has happened, which generates the genotypes $A_2/A_2$, $A_2/A_1$ and $A_1/A_1$ with frequencies ¼, ½ and ¼, respectively. If a meiosis occurs during the sweep, the expected time to its occurrence is given by the mean of a uniform distribution over the interval (0, $t_{s1}$), whose expectation is ½ $t_{s1}$; the probability of this case is $\alpha t_{s1}$. If $t_{mc}$ is the expected time conditioned on no meiotic event during the sweep, we have

$$\alpha^{-1} = \frac{1}{2}\alpha t_{s1}^{2} + (1 - \alpha t_{s1})t_{mc}$$

so that

$$t_{mc} = (\alpha^{-1} - \frac{1}{2}\alpha t_{s1}^{2})(1 - \alpha t_{s1})^{-1} \qquad \text{(S1.4)}$$

$T_{mc} = t_{mc} /(2N)$ is the increase in diversity relative to the ancestral population value during the interval between the completion of the sweep of $A_2/A_1$ and the initiation of the sweep of $A_2/A_2$, conditioned on no meiotic event during the sweep (uppercase letters indicate scaling by 2*N*).

Finally, the time taken for the sweep of $A_2/A_2$ to fixation needs to be considered. If $h > 0$, it can be assumed that $A_1/A_1$ will be outcompeted by the other two genotypes, so that we need only consider the relative frequencies of $A_2/A_2$ and $A_2/A_1$. The expected time to fixation of $A_2/A_2$ can then be found as follows. With weak selection, this case is equivalent to haploid selection with selection coefficient $(1 – h)s$ rather than $hs$. Let the initial frequency of $A_2/A_2$ among the post-meiosis genotypes $A_2A_2$ and $A_2A_1$ be $q_\alpha$ (in the present case $q_\alpha$ = 1/3, but a general value is needed for the case when a meiosis occurs during the sweep – see below).

From standard results ([3] p.757), the time to fixation of $A_2/A_2$ that is contributed by the deterministic phase of the sweep is given by the expression:

$$t_{s2} \approx \frac{1}{(1-h)s} ln(\frac{p_\alpha}{q_\alpha p_2}) \qquad \text{(S1.5a)}$$

where $p_\alpha = 1 - q_\alpha$ and $p_2 = 1 - q_2$ is the frequency of non-$A_2/A_2$ genotypes at the initiation of the final stochastic phase of the sweep.

For the present purpose, the small contributions of the two stochastic phases, at the beginning and end of the sweep (see [3]), will be ignored. From [3] p.758, $p_2 \approx 1/[2N(1 - h)s]$, so that Equation (S1.5a) yields the following expression for the scaled time to fixation of $A_2/A_2$:

$$T_{s2} \approx \frac{1}{2N(1-h)s} \ln\left[\frac{2N(1-h)sp_\alpha}{q_\alpha}\right] \qquad \text{(S1.5b)}$$

With $q_a$ = 1/3, we obtain:

$$T_{s2} \approx \frac{1}{2N(1-h)s} ln[4N(1 - h)s] \qquad \text{(S1.5c)}$$

Putting these results together, the net additional expected diversity relative to the ancestral neutral value after the completion of both sweeps that is contributed by the case when there is no meiotic event during the sweep of $A_2A_1$ is equal to:

$$(1 - \alpha t_{s1})(T_{s1} + T_{s2} + T_{mc}) \qquad \text{(S1.6)}$$

**3. Recovery of diversity during a sweep: a single meiosis during the sweep of $A_2A_1$**

If $\alpha\ t_{s1} < 1$, it can be assumed that at most only a single meiotic event occurs during the sweep of $A_2A_1$ to fixation; as shown above, the probability of such an event is $\alpha\ t_{s1}$. The meiotic event occurs at a time $t_m$ after the start of the sweep of $A_2A_1$, where $t_m$ has the p.d.f. $1/\alpha$ (see Equation 1 of the main text). From standard selection theory, the frequency of $A_2A_1$ at time $t_m$ is given by:

$$\frac{q_m}{p_m} \approx \frac{q_0}{p_0} exp(hst_m) \qquad \text{(S1.7)}$$

where $q_0 \approx 1/(2Nhs)$ is the expected frequency of $A_2A_1$ at the end of the first stochastic phase of the sweep (see [3], p.755).

This expression for $q_m$ can be used instead of $q_\alpha$ in Equation (S1.5a) to obtain $T_{s2}$ for a given value of $t_m$. The expected value of $T_{s2}$ for this case is then given by the following integral:

$$T_{sm2} \approx \frac{\alpha}{2N(1-h)s} \int_0^{\alpha^{-1}} \ln\left(\frac{2N(1-h)sp_m}{q_m}\right) dt_m \qquad \text{(S1.8a)}$$

From Equation (S1.7), we have:

$$\ln\left(\frac{2N(1-h)sp_m}{q_m}\right) \approx \ln\ [2N(1-h)s]\ +\ ln(\frac{p_0}{q_0})\ -\ hst_m$$

Substituting this expression into Equation (S1.8a), approximating $p_0$ by 1 and $q_0$ by $1/(2Nhs)$, and integrating, we obtain:

$$T_{sm2} \approx \frac{1}{2N(1-h)s}\{\ln\ (2N(1-h)s)\ +\ \ln\ (2Nhs)\ -\frac{1}{2}\alpha^{-1}hs\} \qquad \text{(S1.8b)}$$

In this case, however, there is no contribution from $T_{s1}$ to the final expression, since the second sweep is initiated by the meiotic event, whose conditional expected waiting time is $\frac{1}{2}t_{s1}$ (see paragraph below Equation S1.3b). The net extra relative diversity contribution from this case (which has probability $\alpha t_{s1}$) is thus:

$$\alpha t_{s1}T_{sm2} + \frac{1}{2}\alpha t_{s1}^2 \qquad \text{(S1.9)}$$

Using Equation (S1.4), the net expected relative post-sweep diversity (where $N_e \approx N$ is the effective size that determines the level of neutral variability in the pre-sweep population) is thus:

$$\frac{\pi}{\pi_0} \approx 2r(1-r) + (2N_e\alpha)^{-1} + \alpha t_{s1}T_{sm2} + (1-\alpha t_{s1})(T_{s1} + T_{s2}) \qquad \text{(S1.10a)}$$

This can be rewritten in terms of scaled parameters as:

$$\frac{\pi}{\pi_0} \approx 2r(1-r) + (2N_e\alpha)^{-1} + 2N_e\alpha T_{s1}T_{sm2} + (1-2N_e\alpha T_{s1})(T_{s1} + T_{s2}) \qquad \text{(S1.10b)}$$

This expression is likely to be dominated by the first two terms, given the requirement that the duration of the sweep of $A_2A_1$ has to be sufficiently short that at most a single meiotic event can occur.

**4. Model of a sweep of a heterozygote for a beneficial mutation when meiosis is rare: inclusion of inbreeding**

The above results can be extended to allow for inbreeding, such that zygotes are not produced by random mating, but instead there is probability of $F_{IS}$ that a pair of uniting gametes are i.b.d. (Wright's fixation index [4]). As before, we denote the initial state of a sweeping diploid individual as $A_2B_2/A_1B_1$, but now $B_1$ and $B_2$ have a probability $1 - F_{IS}$ of non-i.b.d. In the absence of a meiosis during the sweep, $A_2B_2/A_1B_1$ proceeds to fixation, and produces $A_2B_2/A_1B_1$ gametes in the same proportions as before, once a meiosis occurs. In this case, however, $B_1$ and $B_2$ are i.b.d. with probability $F_{IS}$, so that the net probability of non-i.b.d. at the B locus is $2r(1 - r)( 1 - F_{IS})$. Neutral diversity $\pi_0$ in the pre-sweep population is reduced by the factor $1/(1 + F_{IS})$ compared with the outcrossing case, since inbreeding causes the pre-sweep effective population to be reduced from $N$ (if a Wright-Fisher population of size $N$ is assumed, as in the simulations) to $N_e = N/(1 + F_{IS})$ [5]. Similar considerations apply to the case of a single meiosis during the sweep, producing a final probability of non-i.b.d. of $2r(1 - r)(1 - F_{IS})$.

The recovery of diversity can be modelled in the same way as before, except that $N_e = N/(1 + F_{IS})$ [5] should be used for $\pi_0$ and for the various $T$'s. We thus obtain the final expression:

$$\frac{\pi}{\pi_0} \approx 2r(1 - r)(1 - F_{IS}) + (2N_e\alpha)^{-1} + 2N_e\alpha T_{s1} T_{sm2} + (1 - 2N_e\alpha T_{s1})(T_{s1} + T_{s2}) \quad \text{(S1.11)}$$

Exact treatments of two or more meioses per sweep are, as before, very laborious and will not be pursued here. With numerous meioses per sweep, the simulation results with outcrossing suggest that $r$ can be replaced by an effective recombination $r\alpha$, and that this can be used in the standard sweep formulae to calculate $\pi/\pi_0$ [3]. This assumption should also be applicable here, with $r\alpha$ being replaced by $r\alpha(1 - F_{IS})$ to a good level of approximation [3,6].

The selection equation for a given $s$ and $h$ used in the sweep formulae approximation needs to be modified when there is inbreeding, with $h$ being replaced with $F_{IS} + h(1 - F_{IS})$ and $(1 - 2h)$ with $(1 - 2h)(1 - F_{IS})$ [3]. In the asexual phase, $2hs$ should be replaced with $2[F_{IS} + (1 - F_{IS})h]s$. These substitutions should be valid provided that $s^2$ can be neglected compared with $s$.

The net result of these changes is that the effective recombination rate is reduced by inbreeding, approaching zero as $F_{IS}$ approaches 1, and the fixation times are also reduced (c.f. Fig. 2 of [3]), greatly enhancing the effect of a sweep compared with the outbreeding case.

**5. Model of a sweep in an organism where the asexual phase is haploid**

Here, we assume that there is a mating type locus that enforces strict outcrossing. Meiosis is assumed to follow zygote formation, and the haploid products undergo numerous mitotic divisions before a subsequent burst of meioses occurs among all individuals in the population. In this case, we can assume that a mutation to $A_2$ from $A_1$ occurs at a random time during the asexual phase and produces a haploid genotype $A_2B_2$, where $B_2$ denotes an allele at a neutral locus. The state of the initial population is denoted by $A_1B_1$, where the expected divergence between $B_1$ and $B_2$, and between the $B_1$'s from two different individuals, is $\pi_0$.

If no meiosis occurs during the sweep, $A_2B_2$ completely replaces $A_1B_1$. If the frequency of meioses is low, the probability of this case is given by the analogue of Equation (S1.1), where $t_{s1}$ is obtained from Equation (S1.3b) with $h$ = 1, *i.e*, the probability is $1 - \alpha t_{s1} = 1 - 2N_e \alpha T_{s1}$. There is no second sweep phase, and the only diversity remaining after the sweep comes from recovery during the sweep, whose value relative to $\pi_0$ is equal to the new value of $T_{s1}$. The net contribution to relative diversity is thus $(1 - 2N_e \alpha T_{s1}) T_{s1}$.

If a meiotic event occurs during the sweep (probability $\alpha t_{s1}$, assuming a low frequency of meiosis), its conditional expected time from the beginning of the sweep is equal to $\frac{1}{2} t_{s1}$ (see Equation S1.9), with $h$ = 1. In the presence of an active mating type locus, $A_2B_2$ individuals can mate only with $A_1B_1$ individuals, creating $A_2B_2/A_1B_1$ zygotes just as before. Assuming that only the $A_2$ haploid products of meiosis become fixed, the probability of non-i.b.d. among random pairs of $A_2$ haploid individuals at the end of the sweep is $2r(1 - r)$. In this case, the net diversity relative to $\pi_0$ that arises during the sweep is equal to:

$$\alpha t_{s1} T_{sm2} + \frac{1}{2} \alpha t_{s1} T_{s1} = 2N_e \alpha T_{s1} (T_{sm2} + \frac{1}{2} T_{s1}) \qquad \text{(S1.12)}$$

where the $T$'s are given by Equations (S1.8b) and (S1.3a) with $h$ = 1.

The net expected diversity at the end of sweep if at most only a single meiosis can occur, expressed in terms of parameters scaled by $2N_e$, is thus given by:

$$\frac{\pi}{\pi_0} \approx T_{s1}\{1 + \left(2N_e\alpha\right)\left[2r(1 - r) + T_{sm2} - \frac{1}{2}T_{s1}\right]\} \qquad \text{(S1.13)}$$

Given that $2N_e\alpha T_{s1} = \alpha t_{s1}$ must be < 1 for this equation to be valid, this result suggests a larger effect of a sweep in reducing diversity than in the diploid case, where $2r(1 – r)$ appears without such a factor (Equation S1.10); this is a consequence of the fact that a sweep with no meiosis leads to fixation without recombination.

With a high frequency of meioses, the results for the diploid case suggest that we can replace $r$ with $r\alpha$ in the version of the standard sweep equation [3], where the expressions for $h$ = ½ is used, but with $s$ being set to twice the value for the diploid case, reducing the duration of the sweep and hence the opportunity for a meioitic event to occur.


[1] Charlesworth B, Charlesworth D. Elements of evolutionary genetics. Roberts and Company; 2010.
