## Supplementary material for "Beyond Recombination: Exploring the Impact of Meiotic Frequency on Genome-wide Genetic Diversity": S2 Appendix

### Model of a sweep with two meiotic events

Here we consider the case when two meioses occur during the sweep of the beneficial mutation $A_2$ to fixation, using the basic model described in the Appendix to the main text. In this case, all the genotypes carrying $A_2$ that are produced by the first meiotic event must be considered, and their frequencies at the time of the second event determined. We assume that the frequency of meiosis ($\alpha$) is sufficiently high that the first meiosis occurs before the fixation of $A_2A_1$ in the initial sweep.

The products of the first meiosis can be divided into two categories. Category 1 is the cross between $A_2B_2/A_1B_1$ and $A_1B_1/A_1B_1$; category 2 is the cross among $A_2B_2/A_1B_1$ individuals. If the frequency of $A_2B_2/A_1B_1$ at the time of the first meiosis is $q_1$ and $p_1 = 1 - q_1$, the frequencies of these two crosses among those that generate individuals carrying $A_2$ are $P_0 = p_1q_1/(1 - p_1^2)$ and $Q_0 = q_1^2/(1 - p_1^2)$, respectively.

The frequencies of the genotypes carrying $A_2$ that are generated from these two crosses (conditioned on the frequencies of the crosses) are as follows, where $B_3$ denotes an allele at the B locus that is sampled randomly from the $A_1A_1$ population. All $B_3$ alleles are assumed to be non-i.b.d. with each other and with $B_1$ and $B_2$, which are also mutually non-i.b.d.

Category 1

1.1 $\frac{1}{2}(1-r)$ $A_2B_2/A_1B_3$ 1.2 $\frac{1}{2}r$ $A_2B_1/A_1B_3$

Category 2

2.1 $\frac{1}{4}(1-r)^2$ $A_2B_2/A_2B_2$ 2.2 $\frac{1}{4}r^2$ $A_2B_1/A_2B_1$

2.3 $\frac{1}{2}(1-r)^2$ $A_2B_2/A_1B_1$ 2.4 $\frac{1}{2}r(1-r)$ $A_2B_2/A_2B_1$

2.5 $\frac{1}{2}r(1-r)$ $A_2B_2/A_1B_2$ 2.6 $\frac{1}{2}r(1-r)$ $A_2B_1/A_1B_1$

2.7 $\frac{1}{2}r^2$ $A_2B_1/A_1B_2$

As expected, the sum of the frequencies for category 2 is ¾, the fraction of $A_2$ carrying individuals in an F2 cross.

The expected time to the first meiosis is $t_1 = \frac{1}{2}\alpha^{-1}$ (Equation A2 of the Appendix). As described in the Appendix, the selection process during the initial sweep is that for asexual lineages; here, $A_2A_1$ has a selective advantage *hs* over $A_1A_1$. Using Equation (A7), the frequency of $A_2A_1$ at this time is given by:

$$\frac{q_1}{p_1} \approx \frac{q_0}{p_0} exp(hst_1) \qquad (S1)$$

where $q_0 \approx 1/(2Nhs)$ is the expected frequency of $A_2A_1$ at the end of the first stochastic phase of the sweep. This expression will be used as an approximation for determining the subsequent behaviour of the system, ignoring the stochastic element in the timing of the first meiosis.

The frequency of $A_2A_1$ individuals among the progenies of the two categories of crosses, conditioned on the presence of $A_2$, is given by:

$$P = \frac{1}{2}(P_0 + Q_0)/(\frac{1}{2}P_0 + \frac{3}{4}Q_0) = (1 - \frac{1}{2}q_1)/(1 - \frac{1}{4}q_1) \qquad (S2)$$

and the corresponding frequency of $A_2A_2$ individuals is

$$Q = 1 - P = (1 - \frac{1}{4}q_1)/(1 - \frac{1}{4}q_1) \qquad (S3)$$

The expected time to the second meiosis following the first meiosis is $t_2 = \alpha^{-1}$. After the first meiosis, the relative frequencies of $A_2A_1$ and $A_2A_2$ individuals change according to the equation for an asexual variant with a relative fitness advantage to $A_2A_2$ over $A_2A_1$ of $(1 + s)/(1 + hs) \approx 1 + (1 - h)s$. After $t_2$ generations, the ratio $Q/P$ will thus become equal to:

$$R = \frac{Q}{P}\left(\frac{1+s}{1+hs}\right)^{t_2} \approx \frac{Q}{P}\exp\ [(1-h)\ st_2] \qquad \text{(S4)}$$

and the conditional frequencies of $A_2A_1$ and $A_2A_2$ become:

$$P' = \frac{1}{1+R'}\ ,\ Q' = \frac{R'}{1+R'} \qquad \text{(S5)}$$

The relative frequencies of the different classes of $A_2A_1$ individuals among all $A_2A_1$ individuals in the above list are unchanged by selection, and similarly for $A_2A_2$. However, the frequencies of $A_2A_2$ relative to $A_2A_1$ among these progenies will be modulated by the factor $R'$. Given these relative frequencies, it is then possible to evaluate the relative frequencies of all possible $A_2A_2$ genotypes in the crosses between the progenies of the two categories of crosses involved in the first meiotic event, which are shown in the table below.

The results can be used to obtain the final state of the population when $A_2A_2$ has become fixed. With random mating, we only need to know the relative frequencies of the different possible haplotypes ($A_2B_2$, $A_2B_1$ and $A_2B_3$) among individuals carrying $A_2$ alleles derived from both parents (without conditioning on the frequency of $A_2A_2$ progeny), for each type of mating.  If these are summed and the relative frequencies normalised by the resulting sum, the net conditional frequencies of the three haplotypes among $A_2A_2$, $x_1$, $x_2$ and $x_3$ can be found.

The net probability of no i.b.d among random combinations of haplotypes with such frequencies is then given by:

$$p_n = 1 - (x_1^2 + x_2^2) \qquad \text{(S6)}$$

To obtain the net diversity at the B locus relative to the neutral value, the scaled duration of the sweep needs to be considered, as was done for the case of a single meiosis in the Appendix. This requires evaluation of the time from the occurrence of the second meiosis to the fixation of $A_2A_2$, $t_3$. Using a similar argument to that used for Equation (A5b), we have:lii

$$t_3 \approx \frac{1}{(1-h)s} ln[\frac{q_3 P'}{p_3 Q'}] \qquad \text{(S7)}$$

where $p_3 \approx 1/[2N_e(1 - h)s]$ and $q_3 = 1 - p_3$.

Writing $T_i = t_i/(2N_e)$, the net relative diversity is given by:

$$\frac{\pi}{\pi_0} \approx p_n + \sum_{i=1}^{i=3} T_i \qquad \text{(S8)}$$

Here, $\sum_i T_i$ gives the total scaled duration of the sweep of $A_2$ to fixation, including the lag terms reflecting the waiting times to the meiotic events.

If $\alpha \sum_i t_i > 2$, then the expected number of meioses during the sweep must exceed two, and this model is inapplicable. In addition, if $t_3 < 0$ in Equation (S7), $A_2$A2 must have become fixed before the second meiosis, and the model is again inpapplicable. There is, therefore, only a narrow range of *α* values of a for a given selection regime to which this model can be applied.

### Table of mating frequencies and resulting haplotype frequencies

Here, $P_1 = P$, $Q_1 = Q$ and $Q_2 = RQ$.

| Mating frequency | | Haplotype frequencies | | | $A_2A_2$ |
|---|---|---|---|---|---|
| | | $A_2B_2$ | $A_2B_1$ | $A_2B_3$ | (check) |
| 1.1 x 1.1 | ¼ $P_1^2(1 - r)^2$ | ¼ $(1 - r)$ | 0 | ¼ $r$ | ¼ |
| 1.1 x 1.2 | ½ $P_1^2 r(1 - r)$ | (1/8) $(1 - r)$ | (1/8) $(1 - r)$ | ¼$r$ | ¼ |
| 1.2 x 1.2 | ¼ $P_1^2 r^2$ | 0 | ¼ $(1 - r)$ | ¼$r$ | ¼ |
| 1.1 x 2.1 | ¼ $P_1Q_2(1{-}r)^3$ | ½ $(1 - ½r)$ | 0 | ¼ $r$ | ½ |
| 1.1 x 2.2 | ¼ $P_1Q_2\ r^2(1 - r)$ | ¼$(1 - r)$ | ¼ | ¼ $r$ | ½ |
| 1.1 x 2.3 | ½ $P_1Q_1\ (1 - r)^3$ | ¼$(1 - r)$ | (1/8)$r$ | (1/8)$r$ | ¼ |
| 1.1 x 2.4 | ½ $P_1Q_2 r(1 - r)^2$ | (1/8) $(3 - 2r)$ | (1/8) | ¼ $r$ | ½ |
| 1.1 x 2.5 | ½ $P_1Q_1 r(1 - r)^2$ | (1/8)$(2 - r)$ | 0 | (1/8) $r$ | ¼ |
| 1.1 x 2.6 | ½ $P_1Q_1 r(1 - r)^2$ | (1/8) $(1 - r)$ | (1/8) | (1/8)$r$ | ¼ |
| 1.1 x 2.7 | ½ $P_1Q_1 r^2(1 - r)$ | (1/8) | (1/8)$(1 - r)$ | (1/8) $r$ | ¼ |
| 1.2 x 2.1 | ¼ $P_1Q_2 r(1 - r)^2$ | ¼ | ¼ $(1 - r)$ | ¼ $r$ | ½ |
| 1.2 x 2.2 | ¼ $P_1Q_2 r^3$ | 0 | ¼ $(2 - r)$ | ¼ $r$ | ½ |
| 1.2 x 2.3 | ½ $P_1Q_1 r(1 - r)^2$ | (1/8) $(1 - r)$ | (1/8) | (1/8)$r$ | ¼ |
| 1.2 x 2.4 | ½$P_1Q_2 r^2(1 - r)$ | (1/8) $(3 - 2r)$ | (1/8) | ¼ $r$ | ½ |
| 1.2 x 2.5 | ½ $P_1Q_1 r^2(1 - r)$ | (1/8) | (1/8)$(1 - r)$ | (1/8)$r$ | ¼ |

| | | | | | |
|---|---|---|---|---|---|
| 1.2 x 2.6 | ½ $P_1Q_1r^2(1-r)$ | 0 | (1/8) $(2-r)$ | (1/8)$r$ | ¼ |
| 1.2 x 2.7 | ½ $P_1Q_1r^3$ | (1/8)$r$ | ¼ $(1-r)$ | (1/8)$r$ | ¼ |
| 2.1 x 2.1 | (1/16)$Q_2^2(1-r)^4$ | 1 | 0 | 0 | 1 |
| 2.1 x 2.2 | (1/8)$Q_2^2r^2(1-r)^2$ | ½ | ½ | 0 | 1 |
| 2.1 x 2.3 | ¼ $Q_1Q_2(1-r)^4$ | ¼ $(2-r)$ | ¼ $r$ | 0 | ½ |
| 2.1 x 2.4 | ¼ $Q_2^2r(1-r)^3$ | ¾ | ¼ | 0 | 1 |
| 2.1 x 2.5 | ¼ $Q_1Q_2r(1-r)^3$ | ½ | 0 | 0 | ½ |
| 2.1 x 2.6 | ¼ $Q_1Q_2r(1-r)^3$ | ¼ | ¼ | 0 | ½ |
| 2.1 x 2.7 | ¼ $Q_1Q_2r^2(1-r)^2$ | ¼ $(1+r)$ | ¼ $(1-r)$ | 0 | ½ |
| 2.2 x 2.2 | (1/16) $Q_2^2r^4$ | 0 | 1 | 0 | 1 |
| 2.2 x 2.3 | ¼ $Q_1Q_2r^2(1-r)^2$ | ¼ $(1-r)$ | ¼ $(1+r)$ | 0 | ½ |
| 2.2 x 2.4 | ¼ $Q_2^2r^3(1-r)$ | ¼ | ¾ | 0 | 1 |
| 2.2 x 2.5 | ¼ $Q_1Q_2r^3(1-r)$ | ¼ | ¼ | 0 | ½ |
| 2.2 x 2.6 | ¼ $Q_1Q_2r^3(1-r)$ | 0 | ½ | 0 | ½ |
| 2.2 x 2.7 | ¼ $Q_1Q_2r^4$ | ¼ $r$ | ¼$(2-r)$ | 0 | ½ |
| 2.3 x 2.3 | ¼ $Q_1^2(1-r)^4$ | ¼ $(1-r)$ | ¼ $r$ | 0 | ¼ |
| 2.3 x 2.4 | ½ $Q_1Q_2r(1-r)^3$ | (1/8)$(3-2r)$ | (1/8)$(1+2r)$ | 0 | ½ |
| 2.3 x 2.5 | ½$Q_1^2r(1-r)^3$ | (1/8)$(2-r)$ | (1/8)$r$ | 0 | ¼ |

| | | | | | |
|---|---|---|---|---|---|
| 2.3 x 2.6 | $\frac{1}{2} Q_1^2 r (1 - r)^3$ | (1/8) $(1 - r)$ | (1/8) $(1+ r)$ | 0 | ¼ |
| 2.3 x 2.7 | $\frac{1}{2} Q_1^2 r^2 (1 - r)^2$ | (1/8) | (1/8) | 0 | ¼ |
| 2.4 x 2.4 | $\frac{1}{4} Q_2^2 r^2 (1 - r)^2$ | ½ | ½ | 0 | 1 |
| 2.4 x 2.5 | $\frac{1}{2} Q_1 Q_2 r^2 (1 - r)^2$ | ¼ | ¼ | 0 | ½ |
| 2.4 x 2.6 | $\frac{1}{2} Q_1 Q_2 r^2 (1 - r)^2$ | (1/8) | (3/8) | 0 | ½ |
| 2.4 x 2.7 | $\frac{1}{2} Q_1 Q_2 r^3 (1 - r)$ | (1/8)$(1+2r)$ | (1/8)$(3 - 2r)$ | 0 | ½ |
| 2.5 x 2.5 | $\frac{1}{4} Q_1^2 r^2 (1 - r)^2$ | ¼ | 0 | 0 | ¼ |
| 2.5 x 2.6 | $\frac{1}{2} Q_1^2 r^2 (1 - r)^2$ | (1/8) | (1/8) | 0 | ¼ |
| 2.5 x 2.7 | $\frac{1}{2} Q_1^2 r^3 (1 - r)$ | (1/8)$(1 + r)$ | (1/8)$(1 - r)$ | 0 | ¼ |
| 2.6 x 2.6 | $\frac{1}{4} Q_1^2 r^2 (1 - r)^2$ | 0 | ¼ | 0 | ¼ |
| 2.6 x 2.7 | $\frac{1}{2} Q_1^2 r^3 (1 - r)$ | (1/8)$r$ | (1/8)$(2 - r)$ | 0 | ¼ |
| 2.7 x 2.7 | $\frac{1}{4} Q_1^2 r^4$ | ¼$r$ | ¼ $(1 - r)$ | 0 | ¼ |

The figure below displays results for $\pi/\pi_0$ obtained using these formulae, for the case of a population size of $N$ = 1000, $s$ = 0.1, $h$ = 0.5 and $\rho = 5 \times 10^{-8}$ , with $\alpha$ = 0.0060 (long dashed red curve) and $\alpha$ = 0.0065 (full black curve). For comparison, the results for a single meiosis with $\alpha$ = 0.0040 (short dashed blue curve) are also shown. The curves are all very close to each other, as might be expected.

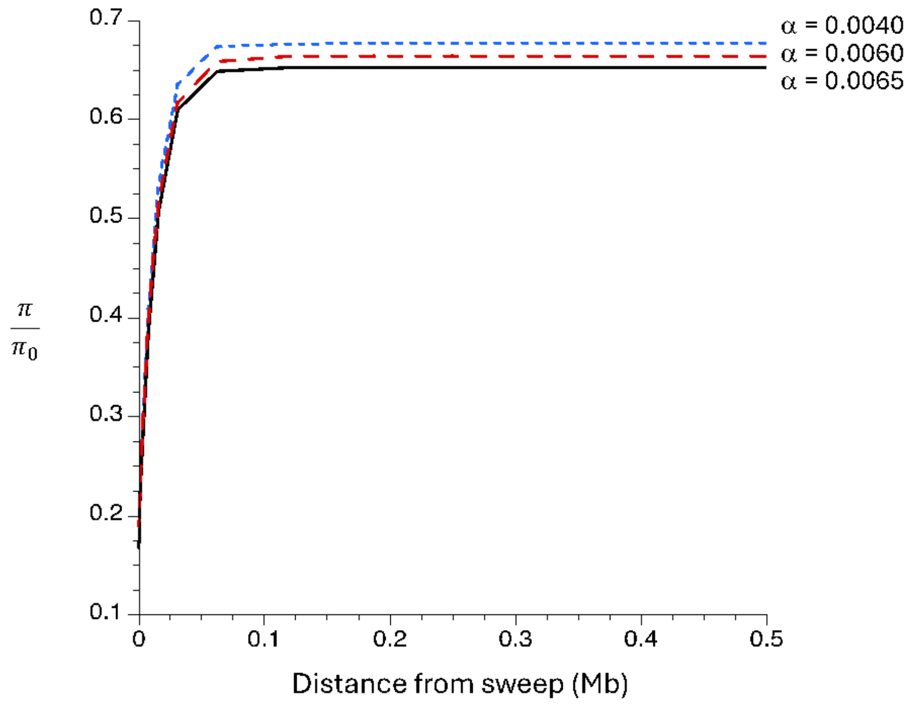
