## Supplementary material for "Beyond Recombination: Exploring the Impact of Meiotic Frequency on Genome-wide Genetic Diversity": S3 Appendix

### Additional analysis of parameters

**1. The effect of the net recombination rate on diversity ………………………………………….... *p. 2***

**2. Recombination rate per meiosis determines the width of the diversity valley……………… *p. 3***

**3. The dominance coefficient slightly affects the loss of genome-wide genetic diversity…… *p. 4***

**4. A high selfing rate causes a significant loss of genome-wide genetic diversity………….... *p. 6***

#### 1. The effect of the net recombination rate on diversity

In canonical models of selective sweeps ($\alpha$ = 1, in our model), the recombination rate per nucleotide site per generation ($\rho$) relative to the strength of selection controls the width of the valley of diversity around the location of the beneficial mutation. We wanted to disentangle the roles played by $\alpha$, $\rho_\alpha = \rho/\alpha$ and $\rho$ (where $\rho_\alpha$ is the recombination rate per sexual generation, and $\alpha$ is the frequency of sexual generations) on genetic diversity, for a given selection regime. We expected to find that $\rho$, the net rate of recombination per generation, controls the width of the valley regardless of $\alpha$, and we also wanted to explore the impact of $\rho_\alpha$ and $\alpha$ on $\pi/\pi_0$ on the unliked chromosome.

Panels A-C in S4 A-C show that, as expected, the smaller $\rho$, the wider the valley around the location of the sweeping mutation on the chromosome, for all $\alpha$ values. In Panel A, the whole chromosome is affected by selection, as recombination is too infrequent to counteract its effect. Neutral loci are in linkage disequilibrium with the beneficial allele until fixation, and genetic diversity does not reach a plateau with increasing distance from the target of selection. In contrast, with a high rate of recombination, diversity reaches a plateau close to its purely neutral value, and the valley is narrower. As expected, diversity on the independent, neutral chromosome (Panel D) is not affected by $\rho$ (no significant difference between each $\rho$ for each $\alpha$, Kruskal-Wallis tests). As is also expected, the time to fixation of the sweep is not affected by $\rho$, taking an average of 425 generations for the beneficial mutation to fix ($p > 0.05$ for all $\alpha$ values, Kruskall-Wallis tests, S8 Fig).

Conversely, for a given $\rho$, $\pi/\pi_0$ on the neutral chromosome decreases with $\alpha$ ($p < 0.05$ for all pairwise $\alpha$ comparisons, except between $\alpha$ = 0.1 and $\alpha$ = 1 for all $\rho$, Dunn's post hoc test). Indeed, the frequency of recombination events between the swept and neutral chromosomes is solely determined by the frequency of sexual reproduction (*i.e*, by $\alpha$ and not by $\rho$) (see S1 Appendix). These results show that $\alpha$ and $s$ both affect the genetic diversity on neutral chromosomes, while $\rho$ affects the width but not the depth of the valley. This is in accord with the predictions of the standard sweep model, given by Equations 14 of [1].

**2. Recombination rate per meiosis determines the width of the diversity valley**

We assumed initially that the effects of sweeps would be controlled by the product $\rho$ of $\alpha$ and the recombination rate per sexual generation $\rho_\alpha$, *i.e*, the recombination rate per generation. We assessed whether this rescaling assumption affects our results and compared the results with a fixed $\rho$ (panel A, same as before) for different $\alpha$ values to a fixed $\rho_\alpha$ (panel B) with the same $\alpha$ values in S5 Fig. As expected, $\pi$ behaves completely differently when $\rho_\alpha$ is not scaled with $\alpha < 1$ (Panel B). Indeed, with $\rho_\alpha = 5 \times 10^{-8}$ and $\alpha < 1$, recombination happens only very rarely. For instance, with $\alpha = 0.01$, $\rho$ equals $5 \times 10^{-10}$ on average per site per generation. With a genome of 2Mbp and approximately 420 generations for the sweep duration (S9 Fig), we have a mean of 0.4 recombination events per individual per simulation ($400 \times 2 \times 10^{6} \times 5 \times 10^{-10}$). The rare recombination events that happen do not effectively break genetic linkage between the beneficial mutation and neutral mutations. This has a strong effect so that, when $\alpha = 0.01$, $\pi/\pi_0$ is ~ 0.2 over the whole of chromosome 1 (Panel B of S5 Fig). As expected given the results from S4 Fig, for a given $\alpha$, $\pi/\pi_0$ on the neutral chromosome is the same on average between the two conditions but the variance of the average $\pi/\pi_0$ is larger when $\rho_\alpha$ is fixed (Panel C of S5 Fig). Once again, the patterns for the mean of $\pi/\pi_0$ are qualitatively consistent with standard sweep theory.

**3. The dominance coefficient slightly affects the loss of genome-wide genetic diversity**

Next, we explore the effect of the dominance coefficient $h$ and $\alpha$ on genetic diversity. So far, we have considered semi-dominant beneficial mutations with $h = 0.5$. We compared $\pi/\pi_0$ for simulations with either partially recessive $h = 0.2$ or partially dominant $h = 0.8$ mutations, see S6 Fig. With $\alpha = 1$ and 0.1, $h$ has no effect on genetic diversity at the plateau nor on the width of the valley (pink line, all panels), in agreement with standard sweep theory [1].

Overall, the genetic diversity ($\pi/\pi_0$) on the neutral chromosome ranges from 0.977 to 1.0 for all $h$ and both $\alpha \geq 0.1$. With $\alpha < 0.1$, the genetic diversity on the second chromosome increases with $h$ (Panel D of S6 Fig). These differences are significant for all three $h$ values with all p-values lower than $8 \times 10^{-4}$ (Dunn's post hoc test). We also compared the simulation results with the theoretical predictions of the standard sweep model (as in Fig 3) for different values of $h$ (S7 Fig). The fit remains good for $\alpha = 1$ for all h values. For $h = 0.2$ and $\alpha \leq 0.1$, the standard model tends to overestimate the diversity on the unlinked chromosome for the lowest $\alpha$ values. The opposite is observed for $h = 0.8$ and $\alpha \leq 0.1$, with the standard model underestimating the diversity on the neutral chromosome with small $\alpha$.

The effects of $\alpha$ and $h$ on the fixation times of beneficial mutations are rather different (S10 Fig). For each $h$, the fixation time is the same between $\alpha = 1$ and $\alpha = 0.1$ ($p > 0.20$, Dunn's post hoc test). If $h = 0.8$, the difference in fixation times is significant when $\alpha < 0.1$. If $h = 0.5$, the difference is significant only when $\alpha = 0.01$ compared to other $\alpha$ values. If $h = 0.2$, the difference is significant between all the pairwise comparisons ($p < 6 \times 10^{-4}$, Dunn's post hoc test, except between $\alpha = 0.01$ and 0.02 as well as $\alpha = 0.1$ and 1).

For each $\alpha$, the fixation time is larger for $h = 0.8$ than for the other two values ($p < 1 \times 10^{-4}$, Dunn's post hoc test), consistent with the slow approach to fixation of dominant mutations in randomly mating populations, due to the sheltering from selection of homozygotes for the recessive alternative allele [1]. The fixation time is larger for $h = 0.2$ than for $h = 0.5$ ($p < 1 \times 10^{-4}$, Dunn's post hoc test, except for $\alpha = 0.01$ where $p = 0.0509$), even though this difference is smaller than to the difference between $h = 0.8$ and 0.5 (< 4% between $h = 0.02$ and $h = 0.05$). These effects of $h$ are predicted by the theory for randomly mating populations (Table 1 of [1]); as in that case, the effects are all small.

#### 4. A high selfing rate causes a significant loss of genome-wide genetic diversity

We investigated the effects of different mating systems on genetic diversity under the standard model ($\alpha$ = 1) and under facultative sex ($\alpha$ < 1), S11 Fig. Genetic recombination during selfing reshuffles alleles within genetically similar backgrounds. We observe a wider valley as the selfing rate increases, regardless of the $\alpha$ value. These results align with simulation results from [2], as well as analytic results in [1] and [2], which show that inbreeding via self-fertilization leads to a wider valley of a reduction in diversity surrounding a selective sweep. The joint effect of selfing and facultative sex results in a stronger reduction of the overall genetic diversity than when only one is acting.

[1] Charlesworth B. How good are predictions of the effects of selective sweeps on levels of neutral diversity? Genetics 2020;216:1217–38. https://doi.org/10.1534/genetics.120.303734.

[2] Hartfield M, Bataillon T. Selective sweeps under dominance and inbreeding. G3 GenesGenomesGenetics 2020;10:1063–75. https://doi.org/10.1534/g3.119.400919.
