## Supplementary material for "Beyond Recombination: Exploring the Impact of Meiotic Frequency on Genome-wide Genetic Diversity": S1 Supplementary figures

**S1 Fig. Times to fixation of the beneficial mutation for different *α* values**
**S2 Fig. The effects of the two models of *α* with hard selective sweeps on genetic diversity (*π*)**
**S3 Fig. Times to fixation of the beneficial mutations under different strengths of selection.**
**S4 Fig. The effect of the recombination rate per basepair on $\pi/\pi_0$, with *s* = 0.05 and *h* = 0.5.**
**S5 Fig. Effect of *ρ* versus $\rho_\alpha$ on $\pi/\pi_0$.**
**S6 Fig. Effects of the dominance coefficient *h*.**
**S7 Fig. The behavior of $\pi/\pi_0$ with different dominance coefficients (*h*) and frequencies of meiosis between theoretical model and simulation**
**S8 Fig. Times to fixation of beneficial mutations under different recombination rates.**
**S9 Fig. Effect of *ρ* versus $\rho_\alpha$ on the time to fixation of a beneficial mutation.**
**S10 Fig. Times to fixation of a beneficial mutation with different dominance coefficients.**
**S11 Fig. Effects of the selfing rate.**

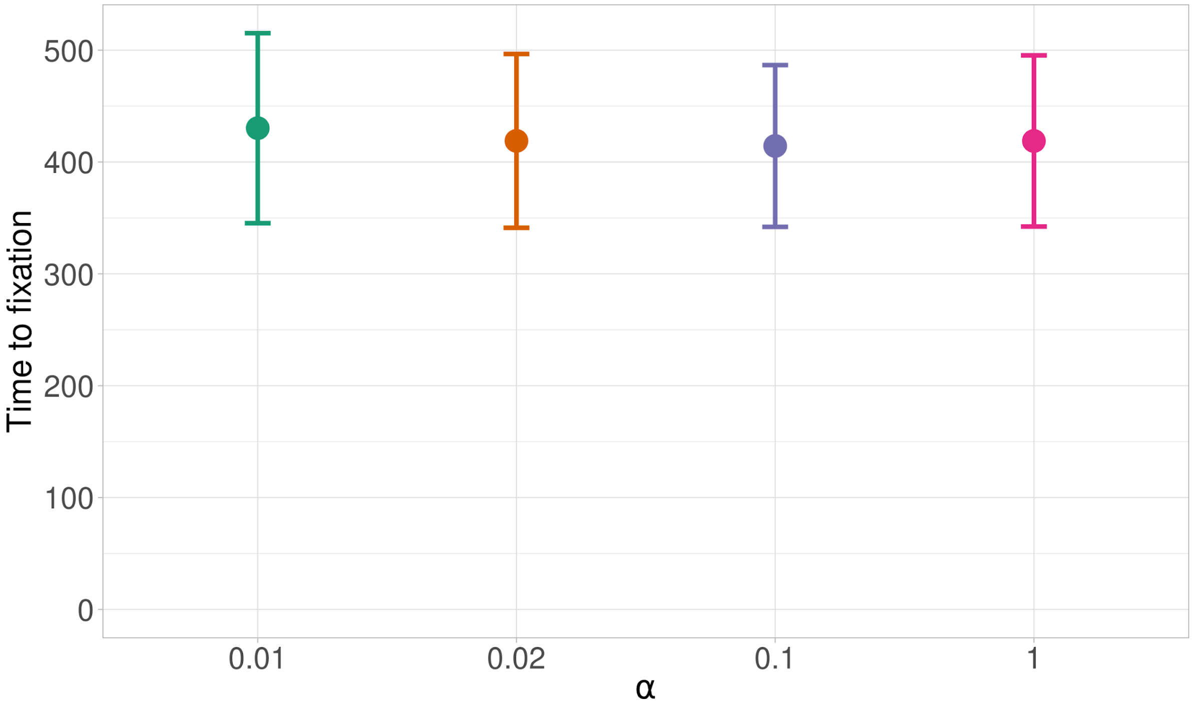


**S1 Fig. Times to fixation of the beneficial mutation for different *α* values.**

Each case used the following settings: a beneficial mutation was introduced into one individual and the population was allowed to evolve until its fixation, with $\rho = 5\times10^{-8}$, $s$ = 0.05 and $h$ = 0.5. If the beneficial mutation was lost, the simulation was rerun. Dots represent the average time to fixation from 500 simulations, with $\alpha$ = 0.01 (green), $\alpha$ = 0.02 (orange), $\alpha$ = 0.1 (blue), and $\alpha$ = 1 (pink). The error bars are standard errors (see Material and Methods for details).

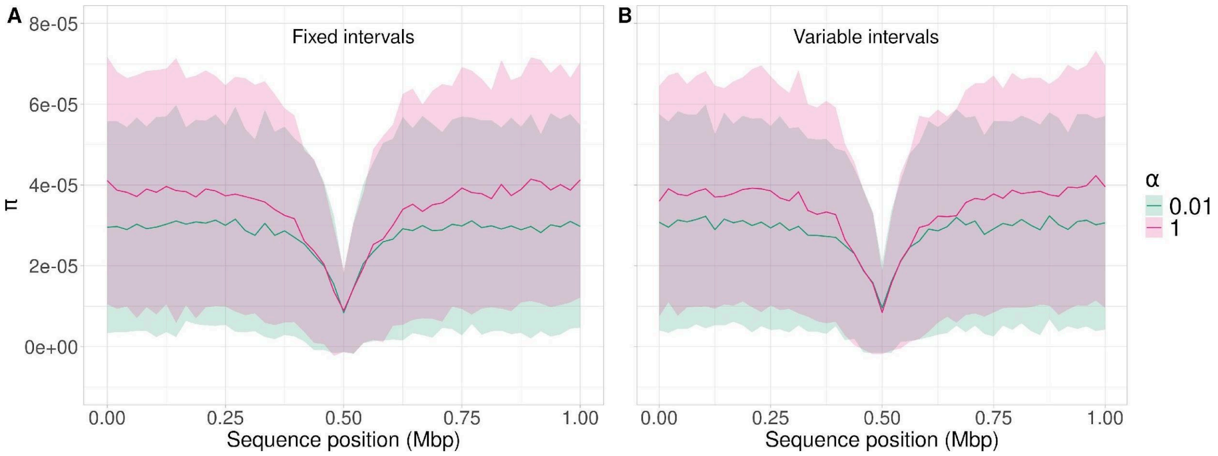


**S2 Fig. The effects of the two models of *α* with hard selective sweeps on genetic diversity (*π)* along a 1 Mb chromosome.**

The same procedure for introducing a beneficial mutation as in S1 Fig was used, with $h$ = 0.5 and $s$ = 0.05. Colors and lines are the same as in Fig 1. With $\alpha$ = 0.01, the interval between two sexual generations was fixed at 1/$\alpha$ = 100 generations (A) or sampled from a Gaussian distribution centered around 1/$\alpha$ = 100 and a standard deviation of 10 (B)

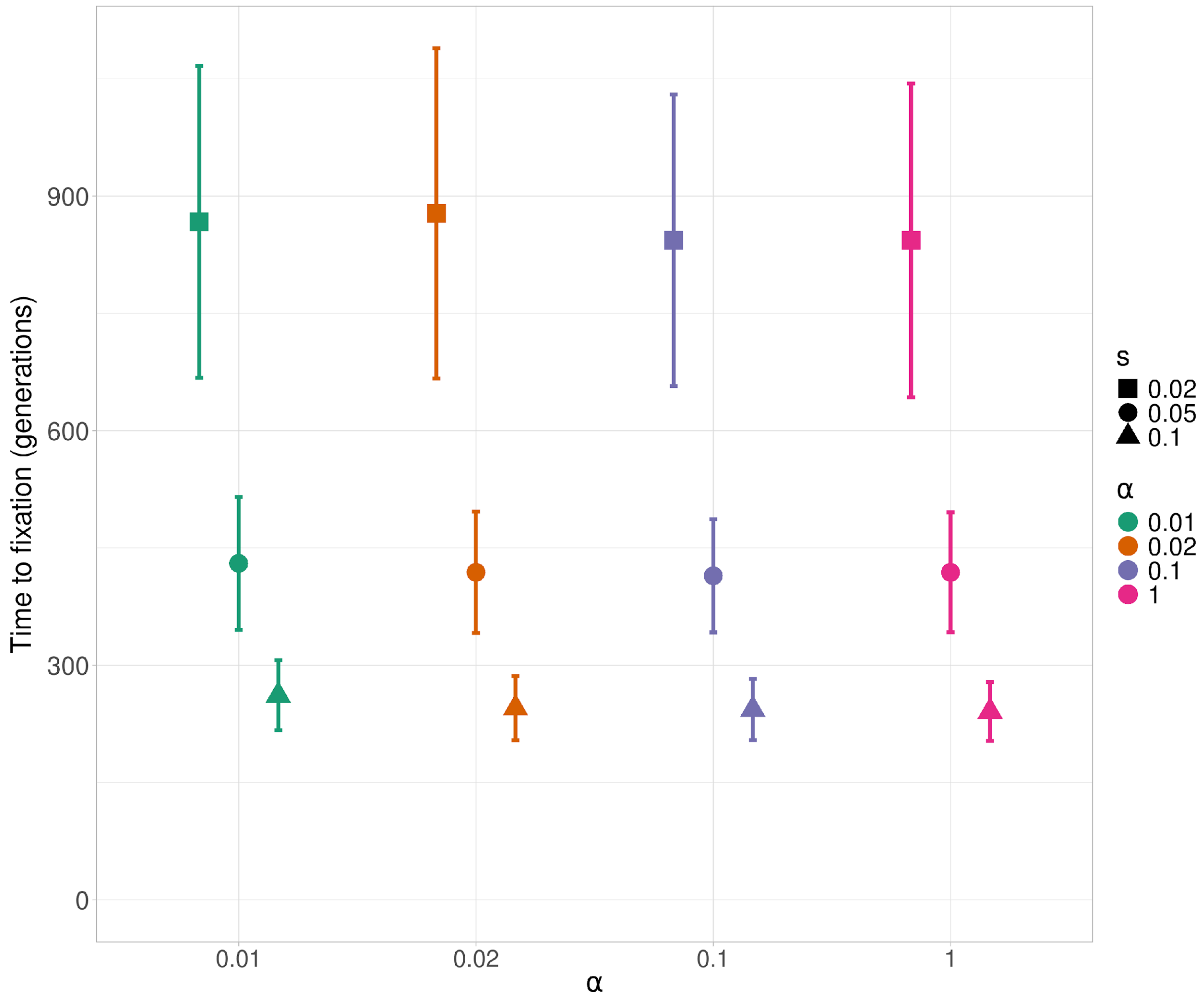


**S3 Fig. Times to fixation of the beneficial mutations under different strengths of selection.**

The same procedure for introducing a beneficial mutation as in the other figures was used, with $\rho = 5 \times 10^{-8}$, $h = 0.5$ and a given $s$. Dots represent the average time to fixation from 500 simulations with $\alpha = 0.01$ (green), $\alpha = 0.02$ (orange), $\alpha = 0.1$ (blue), and $\alpha = 1$ (pink) and error bars are standard errors (see Material and Methods for details).

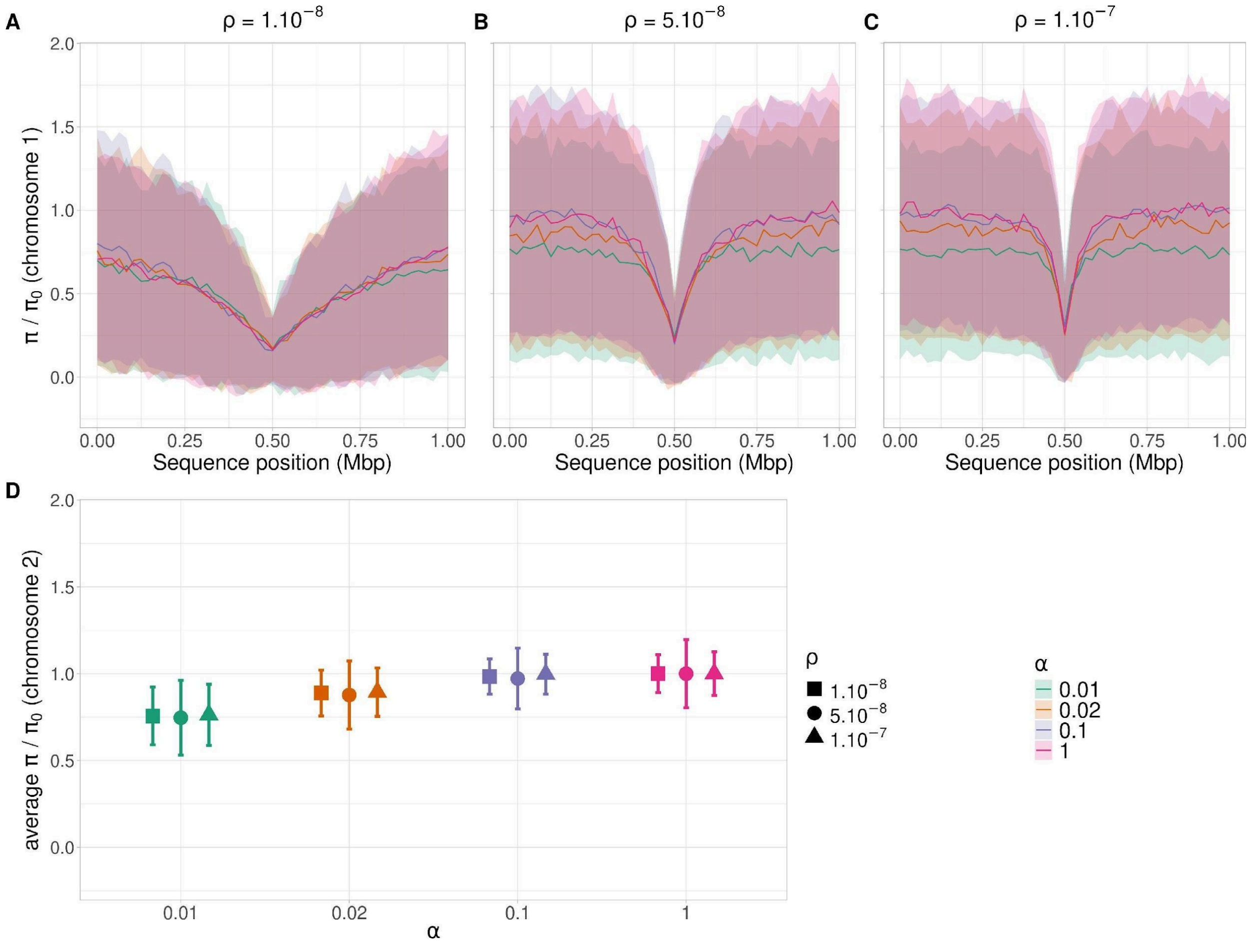


**S4 Fig. The effect of the recombination rate per basepair on $\pi/\pi_0$, with $s$ = 0.05 and $h$ = 0.5.**

(A-C) $\pi/\pi_0$ along a 1Mb chromosome with a hard sweep at 0.5 Mbp. (D) Average $\pi/\pi_0$ and its standard error for a second chromosome that is unlinked to the first one and which carries only neutral mutations. For $\alpha < 1$, the interval between two sexual generations was drawn from a normal distribution centered on $1/\alpha$ generations (see Material and Methods for details). The same procedure for introducing a beneficial mutation as in the other figures was used. The three cases have different recombination rates per generation: $\rho = 1 \times 10^{-8}$ (A), $\rho = 5 \times 10^{-8}$ (B), and $\rho = 1 \times 10^{-7}$ (C). Colored lines are the same as in Fig 2.

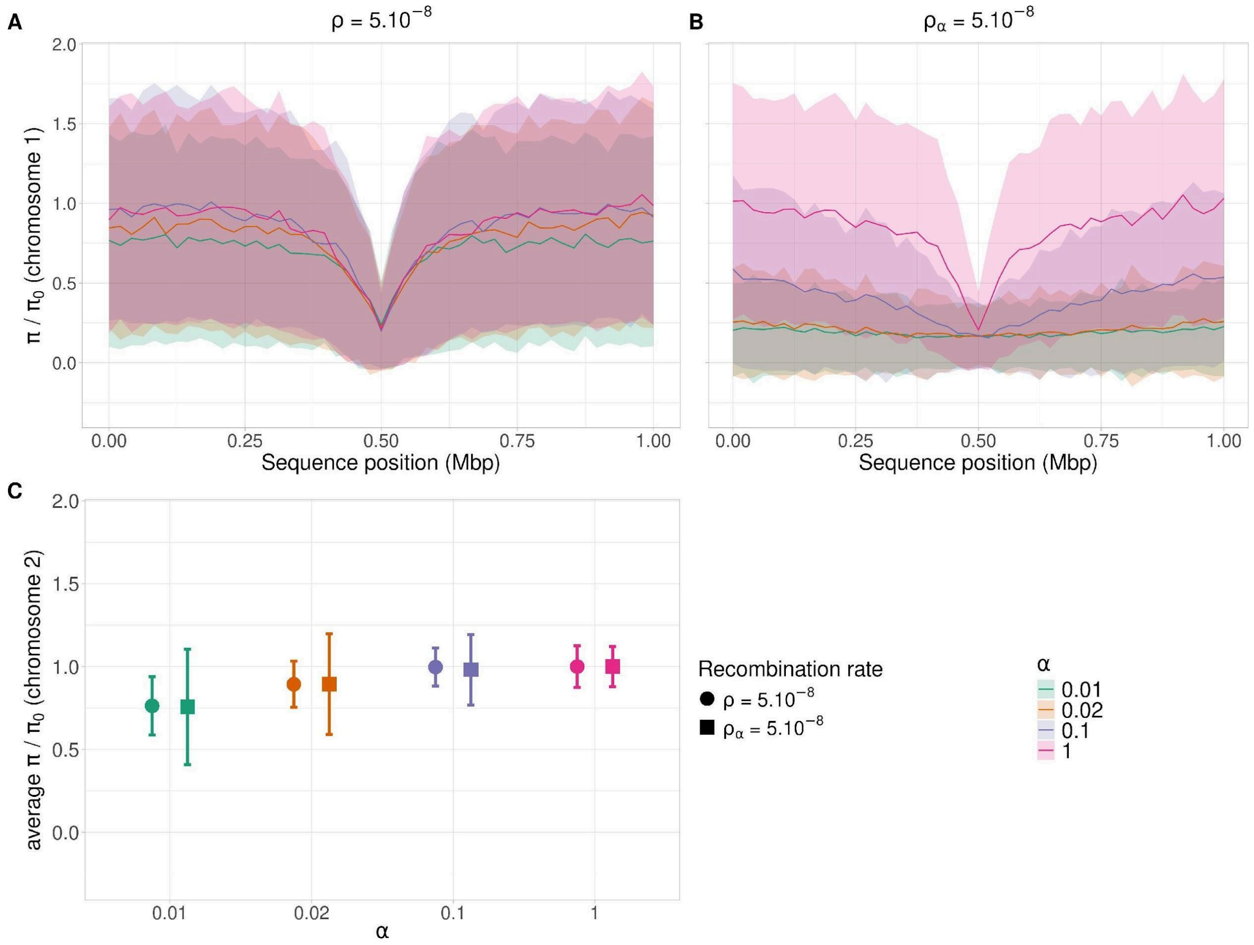


**S5 Fig. Effect of $\rho$ versus $\rho_\alpha$ on $\pi/\pi_0$.**

(A-B): $\pi/\pi_0$ along a 1Mb chromosome with a selective sweep at 0.5 Mbp, with $h$ = 0.5, $s$ = 0.05. (A) Recombination rate per site per generation $\rho = \alpha\, \rho_\alpha$ is constant at $5 \times 10^{-8}$ whereas $\rho_\alpha = 5 \times 10^{-6}$ for $\alpha = 0.01$, $\rho_\alpha = 2.5 \times 10^{-6}$ for $\alpha = 0.02$, $\rho_\alpha = 5 \times 10^{-7}$ for $\alpha = 0.1$ and $\rho_\alpha = 5 \times 10^{-8}$ for $\alpha = 1$. (B) Recombination rate per site per meiosis $\rho_\alpha$ is constant at $5 \times 10^{-8}$ whereas $\rho = 5 \times 10^{-10}$ for $\alpha = 0.01$, $\rho = 1 \times 10^{-9}$ for $\alpha = 0.02$, $\rho = 5 \times 10^{-9}$ for $\alpha = 0.1$ and $\rho = 5 \times 10^{-8}$ for $\alpha = 1$. (C) Average $\pi/\pi_0$ and its standard error for a second, independent chromosome carrying only neutral mutations when $\rho$ is held fixed at the same value (circles) or $\rho_\alpha$ is held fixed (squares). For $\alpha < 1$, the interval between two sexual generations was drawn from a normal distribution centered on $1/\alpha$ generations (see Material and Methods for details). Colored lines are the same as in Fig 2. The same procedure for introducing a beneficial mutation as in other figures was used,

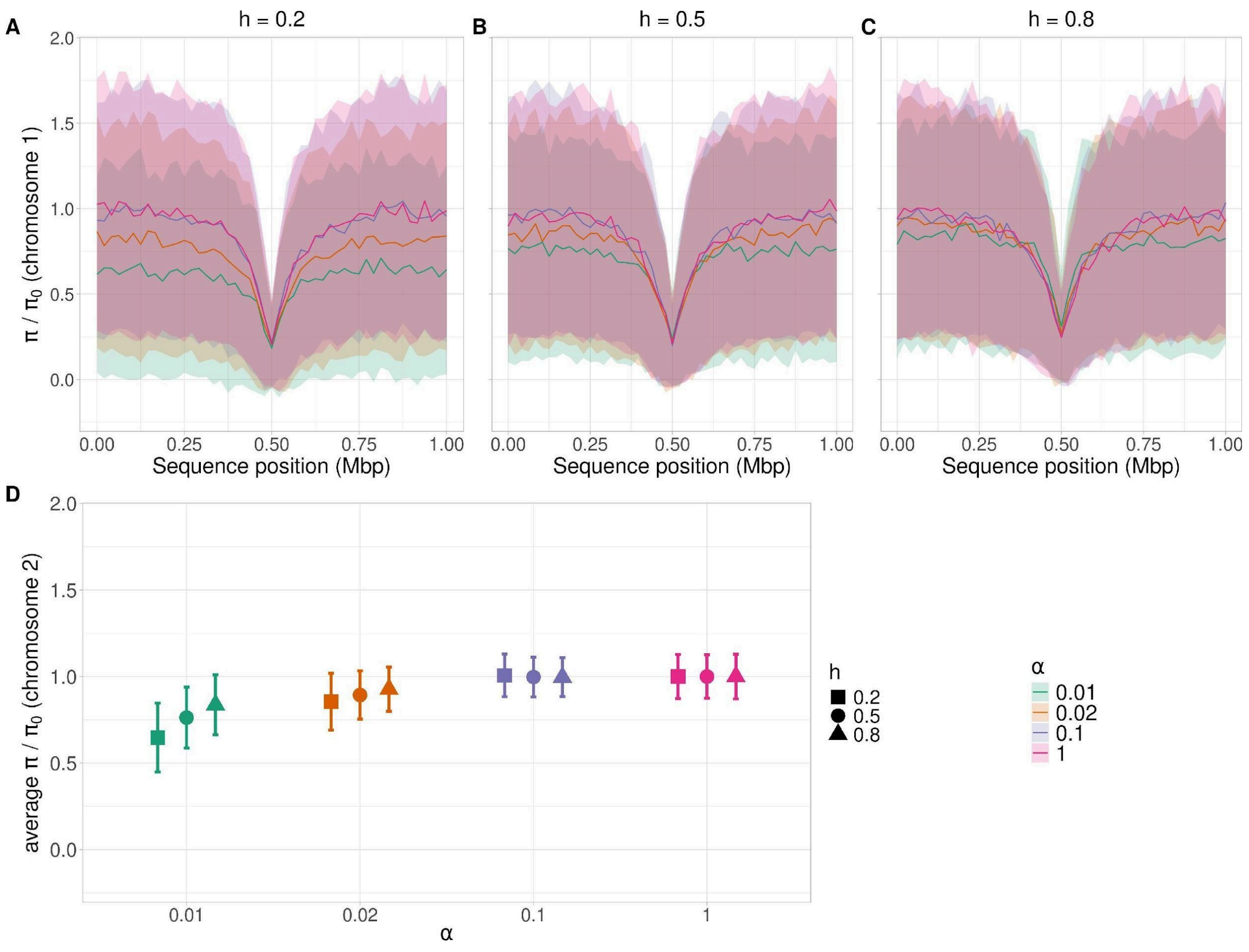


**S6 Fig. Effects of the dominance coefficient *h*.**

(A)-(C) Values of $\pi/\pi_0$ along a 1Mb chromosome that carries a beneficial mutation ($s$ = 0.05) at 0.5 Mbp with $h$ = 0.2 (panel A), 0.5 (B), and 0.8 (C) and $\rho = 5 \times 10^{-8}$. (D) The mean value of $\pi/\pi_0$ (with standard error) on a second, independent chromosome. The same procedure for introducing a beneficial mutation as in other figures was used. For $\alpha < 1$, the interval between two sexual generations was drawn from a normal distribution centered on $1/\alpha$ generations (see Material and Methods for details). Colors and lines are the same as in Fig 1.

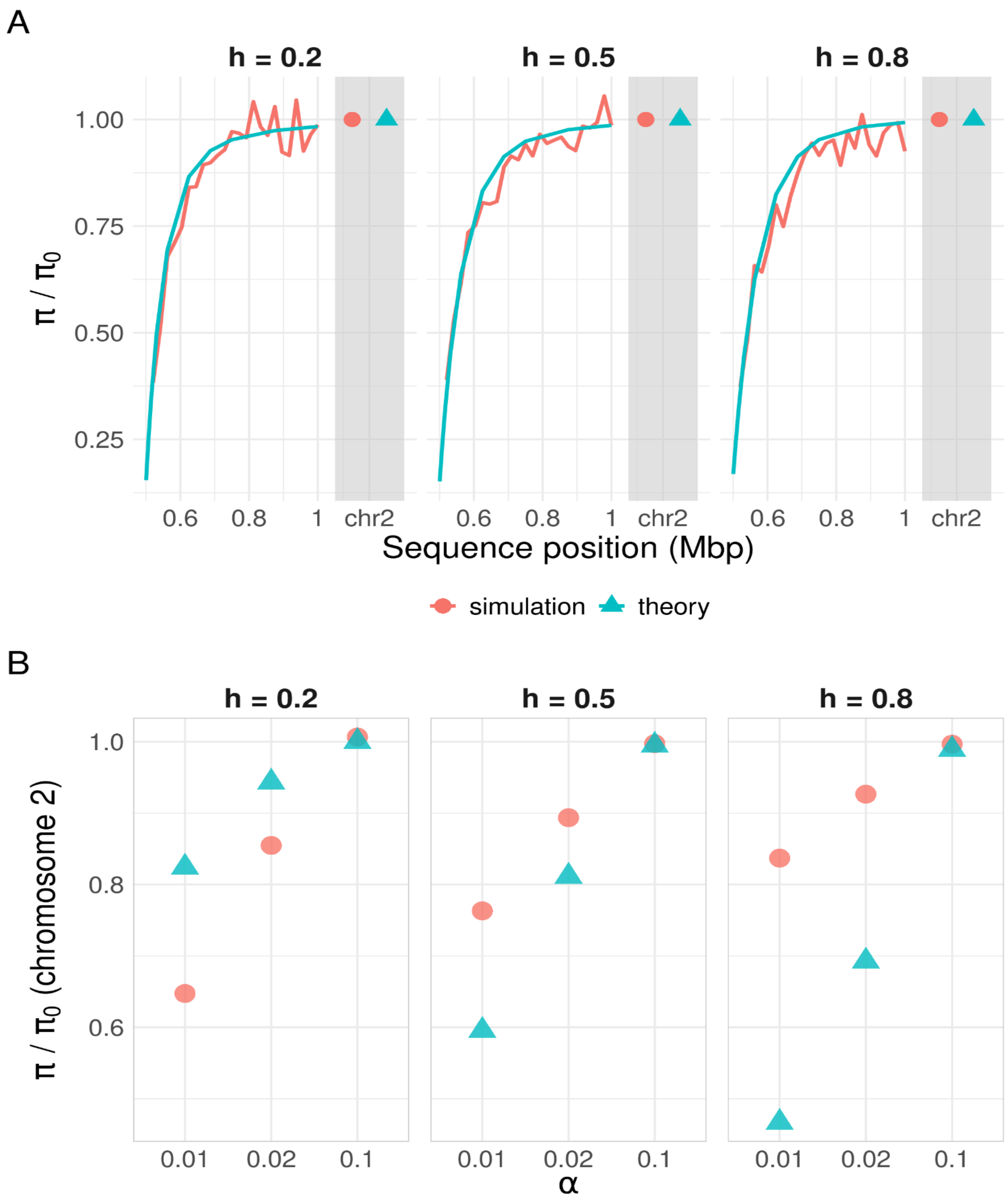


**S7 Fig. The behavior of $\pi/\pi_0$ with different dominance coefficients (*h*) and frequencies of meiosis between theoretical model (Equations 14 from [29], in blue) and simulation results [from Fig 2] in red.**

$\pi/\pi_0$ was computed in the same way as in Fig 2 (see Methods). (A) Comparisons for chromosome 1 (linked) and 2 (unlinked) for $\alpha$ = 1, $s$ = 0.05. (B) Comparisons for chromosome 2, with a combination of $\alpha$ and $h$. Values for simulated data (red) are the averages from 500 simulations. The same procedure for introducing a beneficial mutation as in other figures was used.

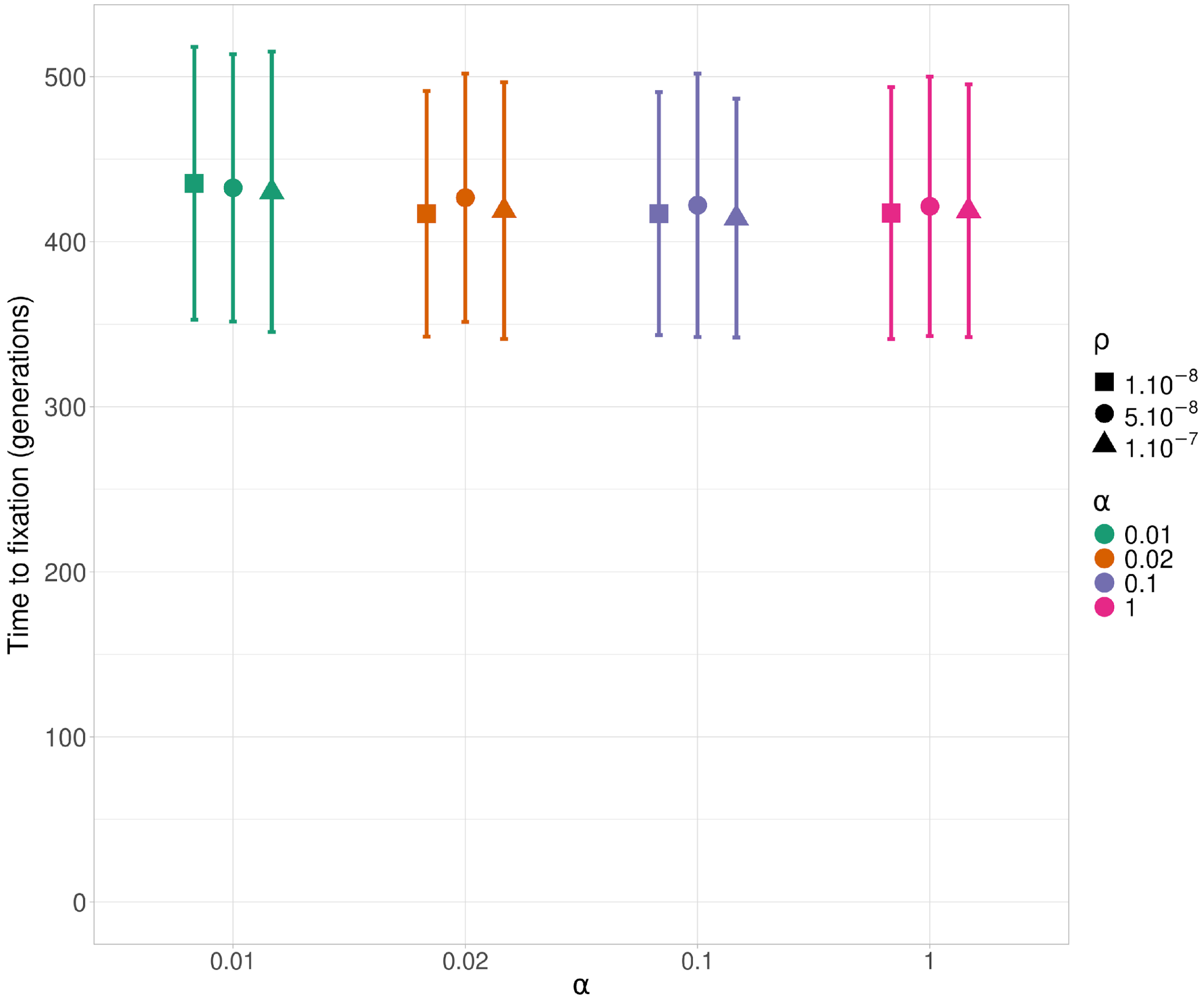


**S8 Fig. Times to fixation of beneficial mutations under different recombination rates**.

The same procedure for introducing a beneficial mutation as in other figures was used, with $h$ = 0.5, $s$ = 0.05 and a given value of $\rho$. Dots represent the average time to fixation from 500 simulations time with $\alpha$ = 0.01 (green), $\alpha$ = 0.02 (orange), $\alpha$ = 0.1 (blue), and $\alpha$ = 1 (pink), and error bars are standard deviation (see. Material and Methods for details).

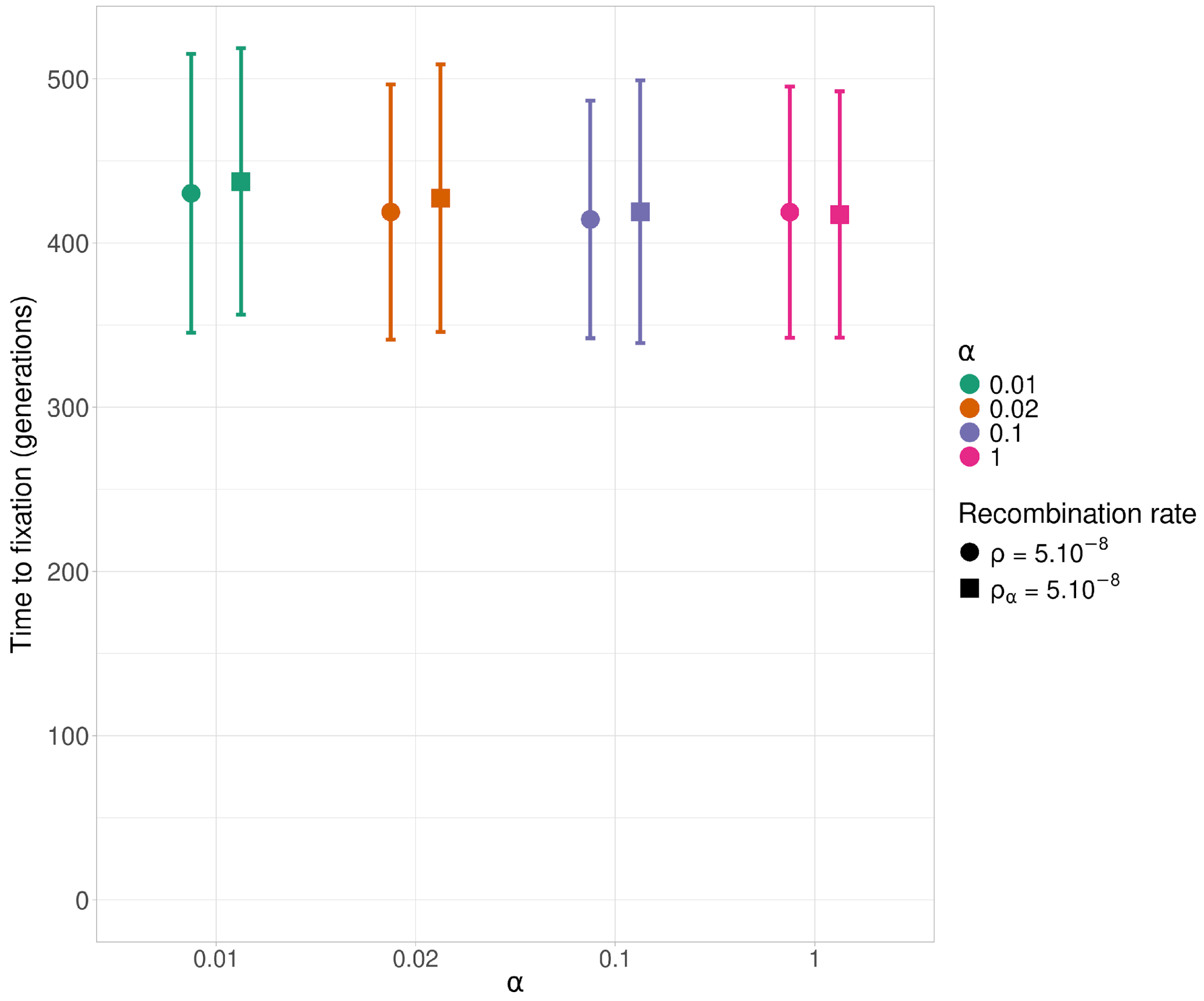


**S9 Fig. Effect of $\rho$ versus $\rho_\alpha$ on the time to fixation of a beneficial mutation.**

The same procedure for introducing a beneficial mutation as in other figures was used, with $h = 0.5$, $s = 0.05$. Either $\rho$ was fixed (circles) or $\rho_\alpha$ was fixed (squares). If recombination rate per site per generation $\rho = \alpha\ \rho_\alpha$ is fixed at $5 \times 10^{-8}$ , we have $\rho_\alpha = 5 \times 10^{-6}$ for $\alpha = 0.01$, $\rho_\alpha = 2.5 \times 10^{-6}$ for $\alpha = 0.02$, $\rho_\alpha = 5 \times 10^{-7}$ for $\alpha = 0.1$ and $\rho_\alpha = 5 \times 10^{-8}$ for $\alpha = 1$. Dots represent the average time to fixation from 500 simulations time with $\alpha = 0.01$ (green), $\alpha = 0.02$ (orange), $\alpha = 0.1$ (blue), and $\alpha = 1$ (pink), and error bars are standard errors (see. Material and Methods for details).

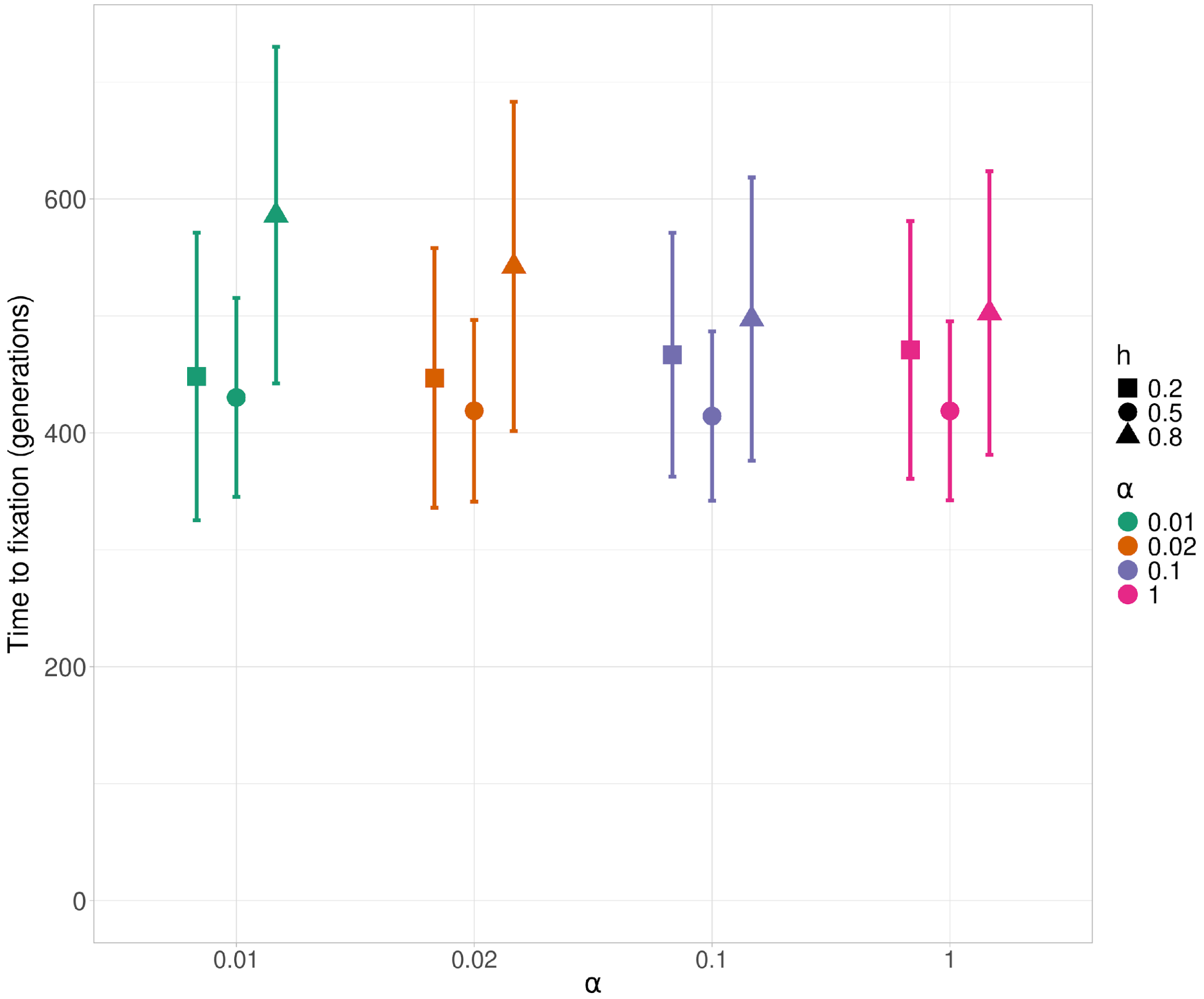


**S10 Fig. Times to fixation of a beneficial mutation with different dominance coefficients.** The same procedure for introducing a beneficial mutation as in other figures was used, with $\rho = 5 \times 10^{-8}$, $s = 0.05$ and a given $h$. Dots represent the average time to fixation from 500 simulations time with $\alpha = 0.01$ (green), $\alpha = 0.02$ (orange), $\alpha = 0.1$ (blue), and $\alpha = 1$ (pink); error bars are standard errors (see. Material and Methods for details).

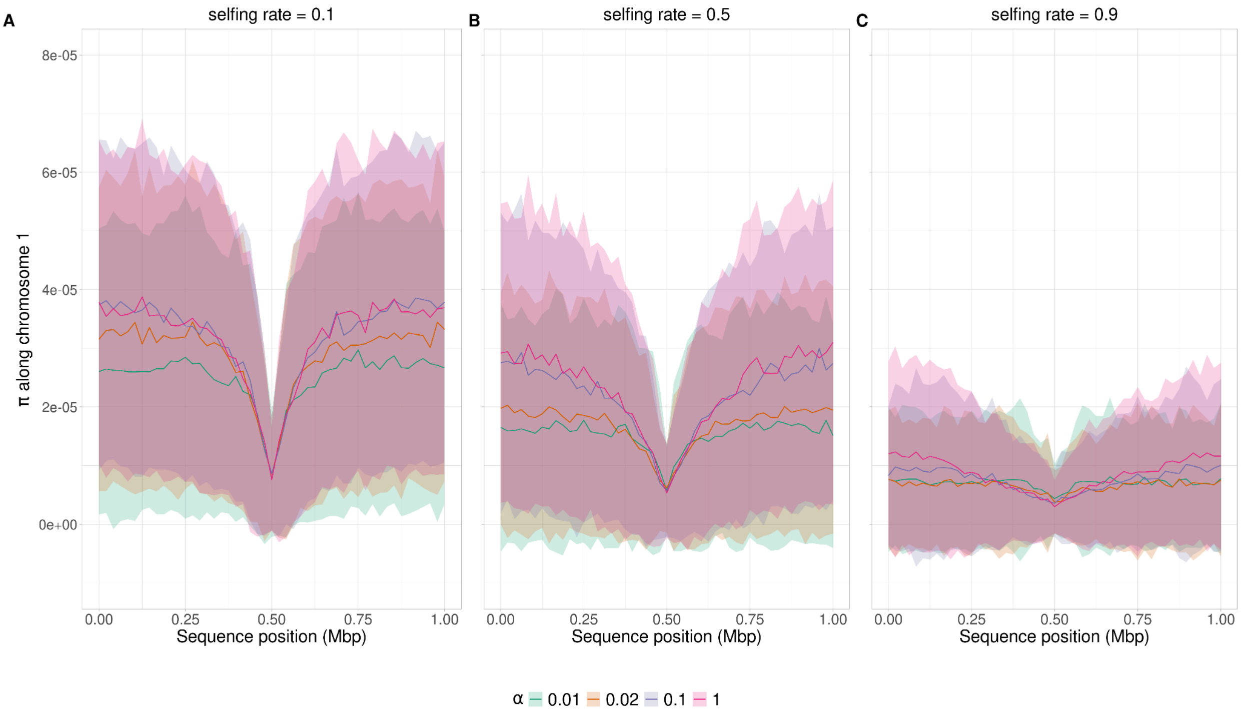


**S11 Fig. Effects of the selfing rate.**

(A)-(C) Values of $\pi$ along a 1Mb chromosome that carries a beneficial mutation ($s$ = 0.05, h = 0.5) at 0.5 Mbp with selfing rate = 0.1 (panel A), 0.5 (B), and 0.9 (C) and $\rho = 5 \times 10^{-8}$. The same procedure for introducing a beneficial mutation as in other figures was used. For $\alpha < 1$, the interval between two sexual generations was drawn from a normal distribution centered on $1/\alpha$ generations (see Material and Methods for details). Colors and lines are the same as in Fig 1.
